# Calcium-independent but voltage-dependent secretion (CiVDS) mediates synaptic transmission in a mammalian central synapse

**DOI:** 10.1101/2021.10.13.464194

**Authors:** Yuan Wang, Rong Huang, Zuying Chai, Changhe Wang, Xingyu Du, Yuqi Hang, Yongxin Xu, Jie Li, Xiaohan Jiang, Xi Wu, Zhongjun Qiao, Yinglin Li, Bing Liu, Xianying Zhang, Peng Cao, Feipeng Zhu, Zhuan Zhou

**Affiliations:** State Key Laboratory of Membrane Biology, Institute of Molecular Medicine, Peking University, Beijing 100871, China; Peking-Tsinghua Center for Life Sciences, Beijing 100871, China; PKU-IDG/McGovern Institute for Brain Research, Beijing 100871, China; Beijing Key Laboratory of Cardiometabolic Molecular Medicine, Beijing 100871, China; National Institute of Biological Sciences, Beijing 102206, China; School of Life Science and Technology, Xi’an Jiaotong University, Xi’an 710049, China

**Keywords:** Ca^2+^-dependent secretion, Ca^2+^-independent but voltage-dependent secretion (CiVDS), synaptic transmission, dorsal root ganglion, dorsal horn, excitatory postsynaptic currents, Cav2.2, SNARE

## Abstract

A central principle of synaptic transmission is that action potential-induced presynaptic neurotransmitter release occurs exclusively *via* Ca^2+^-dependent secretion (CDS). The discovery and mechanistic investigations of Ca^2+^-independent but voltage-dependent secretion (CiVDS) have demonstrated that the action potential *per se* is sufficient to trigger neurotransmission in the somata of primary sensory and sympathetic neurons in mammals. One key question remains, however, whether CiVDS contributes to central synaptic transmission. Here we report, in the central transmission from presynaptic (dorsal root ganglion) to postsynaptic (spinal dorsal horn) neurons, (1) excitatory postsynaptic currents (EPSCs) are mediated by glutamate transmission through both CiVDS (up to 87%) and CDS; (2) CiVDS-EPSCs are independent of extracellular and intracellular Ca^2+^; (3) CiVDS is >100 times faster than CDS in vesicle recycling with much less short-term depression; (4) the fusion machinery of CiVDS includes Cav2.2 (voltage sensor) and SNARE (fusion pore). Together, an essential component of activity-induced EPSCs is mediated by CiVDS in a central synapse.

## Introduction

The tight coupling of synaptic neurotransmitter release to the action potential within a millisecond is critical for neuronal communication and brain function (Augustine, Charlton, & Smith, 1987; Hopfield, 1995; Neher & Sakaba, 2008; Sabatini & Regehr, 1996). The classical view of synaptic transmission is based on the “Ca^2+^ hypothesis”: presynaptic membrane depolarization opens voltage-gated Ca^2+^ channels (VGCCs), which leads to Ca^2+^ influx into the cytosol and triggers vesicular neurotransmitter release (Augustine et al., 1987; Geppert et al., 1994; Jackson & Chapman, 2006; Katz & Miledi, 1967; Neher & Sakaba, 2008; Sudhof, 2012). In this half-century dogma, synaptic transmission is directly triggered by Ca^2+^, synaptotagmin, and SNARE complex assembly (Jahn & Scheller, 2006; Sollner et al., 1993), or membrane depolarization triggers synaptic transmission indirectly by mediating Ca^2+^ influx through VGCCs (Felmy, Neher, & Schneggenburger, 2003; Jackson & Chapman, 2006; Meinrenken, Borst, & Sakmann, 2003; Nanou & Catterall, 2018; Neher & Zucker, 1993; Parnas & Parnas, 2010). Since 2002, this Ca^2+^ hypothesis has encountered exceptions in a series of reports on Ca^2+^-independent but voltage-dependent secretion (CiVDS) in primary sensory dorsal root ganglion (DRG) neurons (Chai et al., 2017; Huang et al., 2019; C. Zhang et al., 2004; C. Zhang & Zhou, 2002; Zheng et al., 2009), where action potentials *per se* directly trigger somatic vesicular exocytosis.

In addition to the DRG (C. Zhang & Zhou, 2002), somatic CiVDS has been extended to trigeminal ganglion neurons (Sforna, Franciolini, & Catacuzzeno, 2019), sympathetic superior cervical ganglion neurons, and neuroendocrine chromaffin cells (Huang et al., 2019; Moya-Diaz et al., 2019). Further studies have revealed unique features of CiVDS compared with Ca^2+^-dependent secretion (CDS): (1) rather than slow and dynamin-dependent endocytosis following CDS (Artalejo, Elhamdani, & Palfrey, 2002; Ferguson et al., 2007; Wu et al., 2019), CiVDS is coupled to a fast and dynamin-independent but protein kinase A-dependent endocytosis (C. Zhang et al., 2004); (2) the vesicle pool replenishment of CiVDS is much faster than that of CDS (Huang et al., 2019; C. Zhang & Zhou, 2002); (3) at the single-vesicle level, CiVDS has a smaller quantal size and faster release kinetics (Huang et al., 2019); and (4) VGCC activation plays dual roles --- indirectly triggering CDS *via* Ca^2+^-influx, and directly triggering CiVDS through the “synprint” binding domain between VGCC and SNARE (Chai et al., 2017; Huang et al., 2019; Nanou & Catterall, 2018). This enables a faster on/off-gating for vesicle fusion *via* CiVDS in somata (Chai et al., 2017; Huang et al., 2019; Liu et al., 2011). The key question remains, however, whether CiVDS occurs in synaptic transmission.

In the present study, by combining electrophysiological recordings, live-cell pHluorin imaging, optogenetic stimulation, and genetic manipulations, we report that CiVDS is present in the central synapses between presynaptic DRG and postsynaptic spinal dorsal horn (DH) neurons. The CiVDS-EPSC has much less short-term depression than that of CDS-EPSC, and predominates during sustained neuronal activity. Altogether, this work provides the first example of CiVDS-mediated synaptic transmission and demonstrates complementary roles of CiVDS and CDS under physiological conditions.

## Results

### CiVDS mediates synaptic transmission in co-cultured DRG and DH neurons

Our previous reports have shown robust CiVDS in the somata of freshly-isolated DRG neurons (Chai et al., 2017; C. Zhang et al., 2004; C. Zhang & Zhou, 2002; Zheng et al., 2009). To investigate whether CiVDS functions in synapses, we co-cultured DRG neurons with postsynaptic DH neurons in which excitatory postsynaptic currents (EPSCs) were collected with whole-cell patch-clamp recordings (***Figure 1A***). Strikingly, local pulses of electrical field stimulation evoked EPSCs in postsynaptic DH neurons, even in 1 mM EGTA (a potent Ca^2+^ chelator)-containing Ca^2+^-free (0Ca) bath solution (***Figure 1A***). Intracellular Ca^2+^ ([Ca^2+^]_i_) recording validated that there was indeed no [Ca^2+^]_i_ rise during recordings in the 0Ca solution (***Figures 1A, Figure1-figure supplement 1***). The amplitude of EPSCs in 0Ca solution was ∼50% of that in the standard 2 mM Ca^2+^ (2Ca) solution (***Figure 1D***). In contrast, the EPSCs recorded in cultured hippocampal neurons were fully dependent on extracellular Ca^2+^ (***Figure 1B***), indicating the absence of CiVDS during synaptic transmission in hippocampal neurons and confirming that the 1 mM EGTA-containing Ca^2+^-free solution was sufficient to block Ca^2+^ transient at synapses.

**Figure 1.**
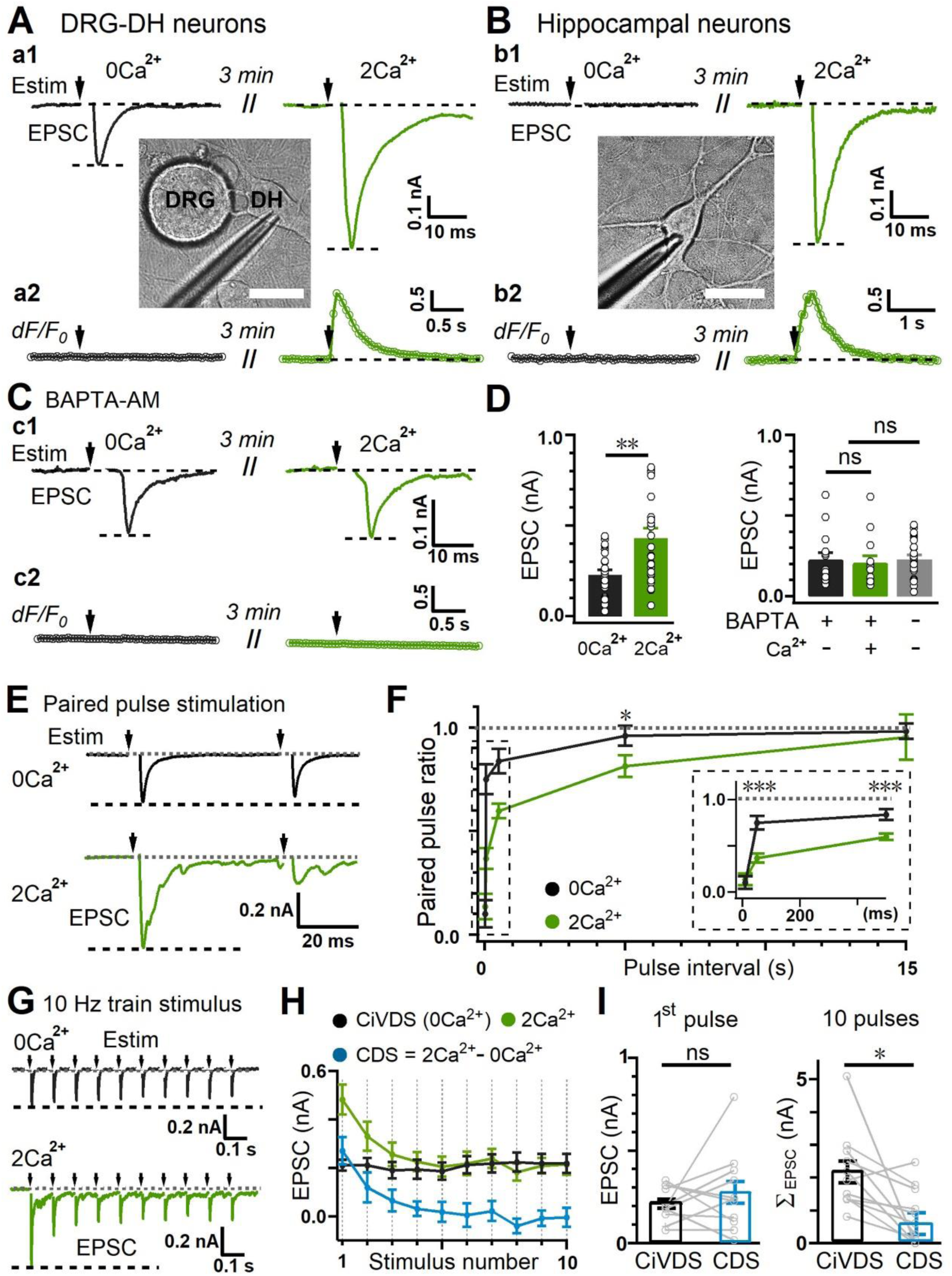
Ca^2+^-independent but voltage-dependent synaptic neurotransmission in co-cultured DRG-DH neurons. (A) (a1) Whole-cell current recording of evoked EPSC signals in response to local electrical field stimulation (Estim, arrows) from a postsynaptic dorsal horn (DH) neuron co-cultured with presynaptic dorsal root ganglion (DRG) neurons in Ca^2+^-free (0Ca^2+^, black) and 2 mM Ca^2+^ bath (2Ca^2+^, green); (a2), evoked intracellular Ca^2+^ signals (dF/F_0_) in a DRG neuron; inset, micrograph showing the setup for EPSC recording from co-cultured DRG-DH neurons (scale bar, 20 μm). (B) (b1) Evoked EPSCs from cultured hippocampal neurons in Ca^2+^-free (black) and 2 mM Ca^2+^-containing solution (green); (b2) evoked intracellular Ca^2+^ signals (dF/F_0_) in hippocampal neurons; inset, micrograph showing the setup for EPSC recordings from a hippocampal neuron (scale bar, 20 μm). (C) As in (A), except that 50 μM BAPTA-AM was pre-loaded into the DRG and DH neurons before recording in Ca^2+^-free (black) or 2 mM Ca^2+^-containing solution (green). (D) Left, quantification of amplitude as in (A) (n = 26 cells for 0Ca^2+^ and 29 cells for 2Ca^2+^). Right, quantification of EPSC amplitudes as in (C) (n = 16 cells for 0Ca^2+^ and 14 cells for 2Ca^2+^). (E) Evoked CiVDS-EPSCs (0Ca^2+^, black) and total EPSCs (CDS + CiVDS, 2Ca^2+^, green) induced by paired-pulse stimulation with a 50-ms interval from DH neurons co-cultured with DRG neurons. The 0Ca^2+^ and 2Ca^2+^ EPSCs were recorded in 2 different DH neurons. (F) Summary plot of paired-pulse ratios as in (E) with different intervals (in 0Ca^2+^, n = 7 cells for 10 ms, 13 for 50 ms, 14 for 500 ms, 10 for 5 s, and 8 for 15 s; in 2Ca^2+^, n = 14 cells for 10 ms, 17 for 50 ms, 22 for 500 ms, 19 for 5 s, and 14 for 15 s). Inset shows the initial plot at an expanded scale. (G) Representative EPSCs induced by a 10-Hz stimulus train in DH neurons co-cultured with DRG neurons in Ca^2+^-free (black) or 2 mM Ca^2+^-containing solution (green). The 0Ca^2+^ and 2Ca^2+^ EPSCs were recorded in 2 different DH neurons. (H) Summary plots of the EPSC amplitudes as in (G), including CiVDS (0Ca^2+^, black), CDS + CiVDS (2Ca^2+^, green), and CDS (2Ca^2+^ – 0Ca^2+^, blue) (n = 11 cells). (I) As in (H), statistics for amplitudes of CiVDS-EPSC (0Ca^2+^, black) and CDS-EPSC (“2Ca^2+^” – “0Ca^2+^”, blue) evoked by single-pulse (first EPSC, left) or 10 pulses (cumulative 10 EPSCs, Σ_EPSC_, right) during 10-Hz train stimulation (n = 11 cells). All but (B) were from DRG and DH co-cultures. Data are shown as the mean ± s.e.m.; Mann-Whitney test for D and F; paired Student’s *t* test for I; *p <0.05, **p <0.01, ***p <0.001, ns, not significant.

To further exclude the possible contribution of intracellular microdomain Ca^2+^, BAPTA-AM, a potent Ca^2+^ chelator, were pre-loaded into the co-cultured neurons. Consistent with our previous reports on somatic secretion (Chai et al., 2017; Huang et al., 2019; C. Zhang & Zhou, 2002), BAPTA had no effect on the EPSCs recorded in the 0Ca solution, but reduced the EPSCs recorded in 2Ca to that in 0Ca (***Figure 1C-D***). Thus, under physiological conditions (2Ca), the evoked EPSC contains both a CiVDS-mediated EPSC (CiVDS-EPSC), and a pure CDS-mediated EPSC (CDS-EPSC), or EPSC (2Ca) = CiVDS-EPSC + CDS-EPSC.

We next assessed the vesicle recycling rate by using paired-pulse stimuli with different time intervals. Consistent with our previous findings in somatic secretion (C. Zhang & Zhou, 2002), the 80% recovery of the total (CiVDS + CDS) EPSC recorded in 2Ca solution required >5 s intervals (***Figure 1E-F***). In contrast, 80% recovery of the CiVDS-EPSC was achieved within 50 ms (***Figure 1E***), indicating a >100 times faster recycling rate/replenishment of vesicle pools in CiVDS than that in CDS in DRG synaptic terminals. In addition, we used physiological 10-Hz train stimulation to evaluate the contribution of CiVDS during sustained synaptic transmission (Fang, McMullan, Lawson, & Djouhri, 2005; Xu, Huang, & Zhao, 2000; Zheng et al., 2009). Intriguingly, CiVDS-EPSCs showed much less short-term depression than the total EPSCs in 2Ca solution (***Figure 1G–H, Figure 1-figure supplement 2***). Importantly, the CiVDS-EPSC contributed 49% of the total EPSC (in amplitude) recorded in the 2Ca bath (CiVDS-EPSC + CDS-EPSC) during single-pulse stimulation, and this increased to 87% during 10-Hz train stimulation (***Figures 1H, Figure 1-figure supplement 2***), suggesting that, in contrast to CDS, CiVDS makes a dominant contribution during sustained neural activity (***Figures 1I, Figure 1-figure supplement 2***).

In addition, we performed live-cell imaging of synaptophysin (Spy)-pHluorin in the nerve terminals of DRG neurons co-cultured with DH neurons. Consistent with EPSC recordings, an electrical field stimulation (20 Hz, 20 s) induced a notable increase of the Spy-pHluorin signal in either 0Ca or 2Ca bath solution (***Figure 2A–C, Figure 2-video 1 and Figure 2-video 2***). Similar to electrophysiological results during 10-Hz stimulation (***Figure 1-figure supplement 2C***), the peak amplitude of ΔF/F_0_ evoked in 0Ca solution was 77% of that evoked in 2Ca solution, further confirming the dominant contribution of CiVDS during sustained neural activity (***Figure 2C***). However, the same electrical stimulation triggered Spy-pHluorin exocytosis in hippocampal nerve terminals only in 2Ca solution, but not 0Ca solution (***Figure 2D–F, Figure 2-video 3 and Figure 2-video 4***). Thus, CiVDS mediates synaptic transmission in co-cultured DRG and DH neurons.

**Figure 2.**
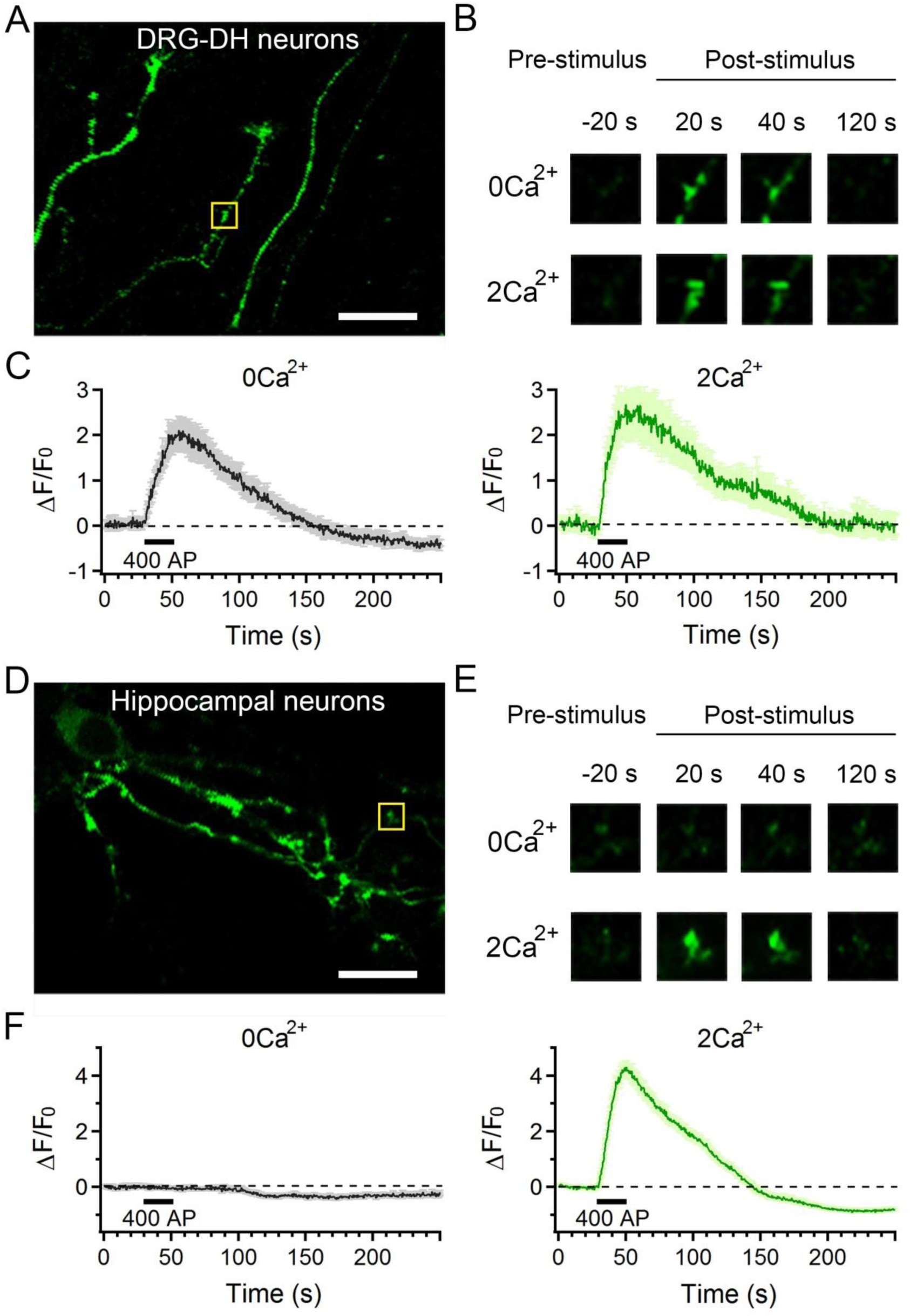
Imaging of CiVDS mediated synaptic transmission in co-cultured DRG and DH neurons. (A) A representative photograph showing the co-cultured DRG and DH neurons. The DRG neurons were expressed with Spy-pHluorin for imaging the synaptic transmission. Scale bar, 10 μm. (B) Images of a presynaptic bouton marked in (A) showing the Spy-pHluorin fluorescence at 20 s before (-20 s, pre-stimulus), 20 s, 40 s, and 120 s after electrical stimulation (post-stimulus) in 0Ca^2+^ (upper panel) or 2Ca^2+^ (lower panel) solution. (C) Averaged fluorescence changes (ΔF/F_0_) of Spy-pHluorin in 0Ca^2+^ (left) or 2Ca^2+^ solution (right) in response to the same electrical stimulation (20 Hz, 20 s) (n = 45 puncta from 6 cells for 0Ca^2+^ and 57 puncta from 6 cells for 2Ca^2+^). The shadow in the traces represents the error bars (s.e.m) of each point. (D-F) The same as in (A-C), but the experiments were performed in cultured hippocampal neurons (n = 70 puncta from 3 cells for 0Ca^2+^ and 72 puncta from 3 cells for 2Ca^2+^).

### CiVDS-EPSC is mediated by glutamate release from presynaptic DRG neurons

To determine whether the CiVDS-EPSC is mediated by presynaptic glutamate release, we tested the effects of ionic glutamate receptor blockers (D-AP5 for NMDA receptor and CNQX for AMPA receptor (Trussell, Zhang, & Raman, 1993; C. Wang et al., 2016)). Three minutes of exposure to AP5 (50 μM) and CNQX (10 μM) diminished the EPSCs in both the 0Ca and 2Ca solutions (***Figure 3A-B***). As a control, there was only minimal run-down during repeated stimulation of CiVDS-EPSCs (***Figure 3-figure supplement 1***). Furthermore, 3-min treatment with 100 μM cyclothiazide (CTZ), a blocker of AMPA receptor desensitization (Fucile, Miledi, & Eusebi, 2006), greatly slowed the decay and increased the charge of the EPSC in both the 0Ca and 2Ca solutions (***Figure 3C-D***). Thus, the CiVDS-EPSC is mediated by synaptic glutamate transmission in co-cultured DRG and DH neurons.

**Figure 3.**
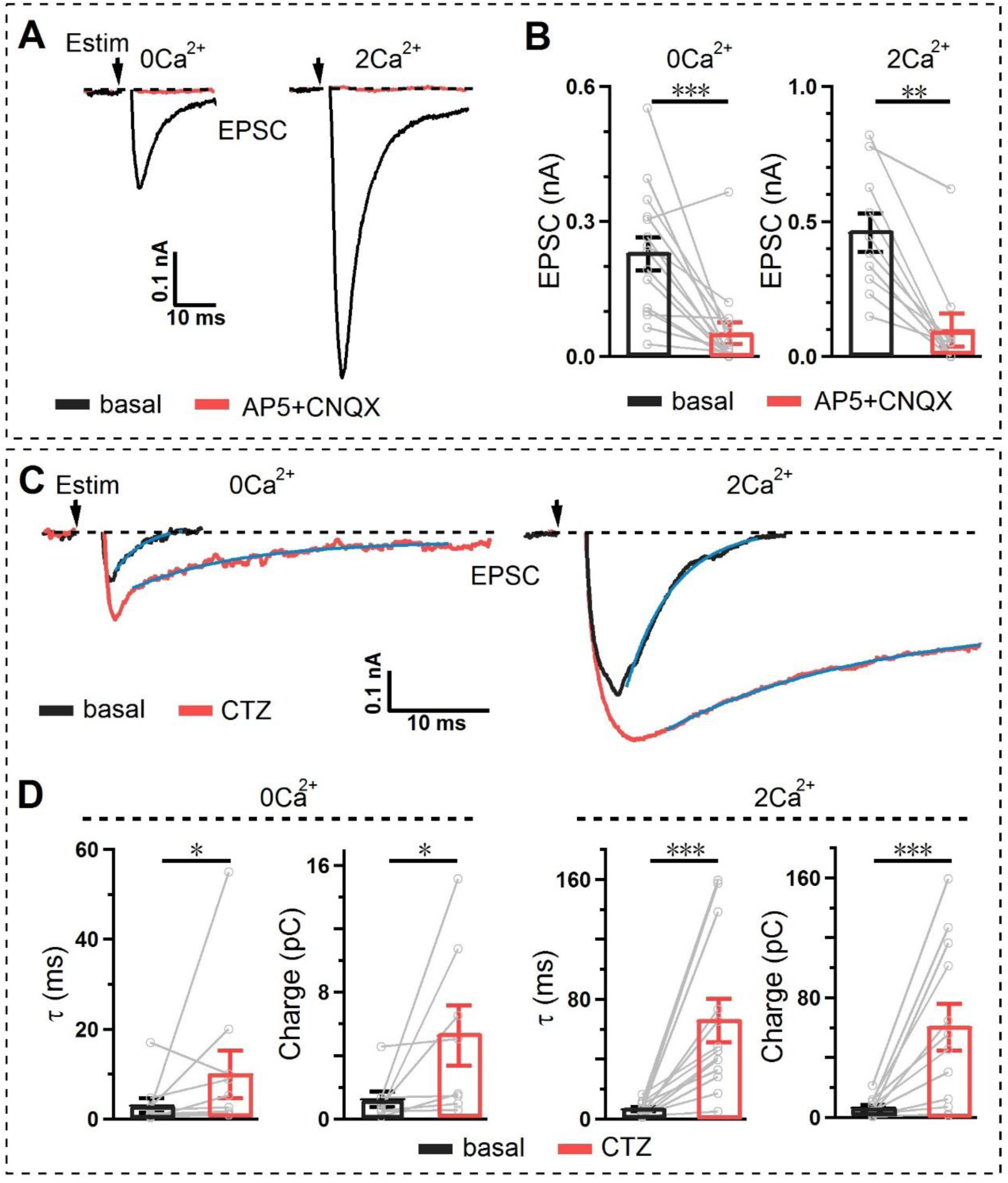
The CiVDS-EPSC is mediated by glutamate transmission. (A) Representative evoked EPSCs in DH neurons co-cultured with DRG neurons before (black) and after (red) applying 50 μM AP-5 and 10 μM CNQX in Ca^2+^-free (left) or 2 mM Ca^2+^-containing solution (right). The EPSCs of 0Ca^2+^ and 2Ca^2+^ were recorded in 2 different DH neurons. (B) Quantification of EPSC amplitudes as in (A) (n = 15 cells for 0Ca^2+^ and 10 cells for 2Ca^2+^). (C) Evoked EPSCs recorded in DH neurons co-cultured with DRG neurons before (black) and after (red) applying 100 μM cyclothiazide (CTZ) in 0Ca^2+^ (left) or 2Ca^2+^ solution (right). The traces are fitted with a single exponential curve (blue). The EPSCs of 0Ca^2+^ and 2Ca^2+^ were recorded in 2 different DH neurons. (D) Quantification of the decay time (τ) and charge as in (C) (n = 10 cells for 0Ca^2+^ and 12 cells for 2Ca^2+^). EPSCs were evoked by local electrical stimulation (Estim) at arrows. Data are shown as the mean ± s.e.m.; paired Student’s *t* test; *p <0.05, **p <0.01, ***p <0.001.

To determine whether DRG neurons are responsible for the presynaptic glutamate release in CiVDS-EPSCs, we performed paired whole-cell recordings to examine the specific synaptic transmission between the patched presynaptic DRG (current-clamp) and postsynaptic DH neurons (voltage-clamp). EPSC traces were recorded in the standard 2Ca bath within 1 min and 0Ca bath 5 min after whole-cell dialysis, in which 10 mM BAPTA was whole-cell dialyzed into the patched DRG neuron (***Figure 4A***) to ensure the intracellular Ca^2+^-free condition (***Figure 4A***). A single pulse of current injection (5 ms, 1000 pA) after break-in triggered an action potential in the presynaptic DRG neuron, followed by an EPSC in the postsynaptic DH neuron in 2Ca bath solution (***Figure 4B***). Strikingly, a notable EPSC signal (∼40%) remained 5 min after whole-cell dialysis even though the patched neurons were bathed in 1 mM EGTA-containing Ca^2+^-free solution (***Figure 4B***). In contrast, similar Ca^2+^-free treatment completely blocked the EPSCs in hippocampal neurons (***Figure 4C-D***). Thus, CiVDS-mediated transmitter release from the presynaptic DRG neurons contributes to EPSC signals in the postsynaptic DH neurons.

**Figure 4.**
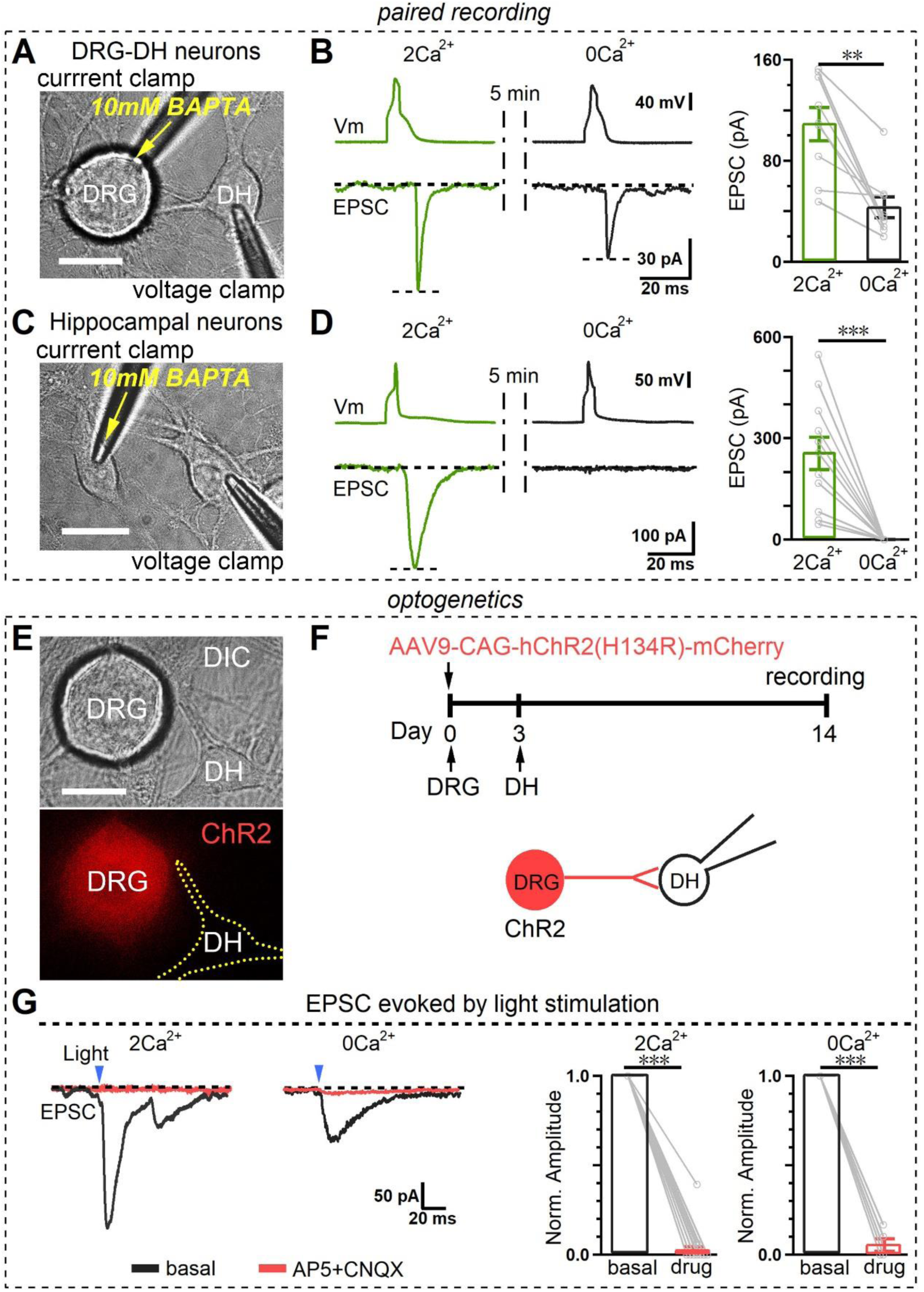
Presynaptic DRG neurons elicit CiVDS-EPSCs in postsynaptic DH neurons. (A) Setup for paired patch-clamp recording of dorsal root ganglion (DRG) and dorsal horn (DH) neurons in co-culture. The presynaptic DRG neuron was whole-cell dialyzed with 10 mM BAPTA under current-clamp mode, while the postsynaptic DH neuron was in voltage-clamp mode. (B) Left, representative dual recordings of presynaptic action potentials (upper) and postsynaptic EPSCs (lower) following current-step injection (1000 pA, 5 ms) in a DRG neuron dialyzed with 10 mM BAPTA. Following whole-cell break-in and intracellular BAPTA dialysis, double recordings were performed at 0 min / 2.5 mM Ca^2+^ bath (green), and 5 min / 0 Ca^2+^ bath (black), Right, statistics of EPSC amplitudes (n = 9 cells). (C and D) As in (A and B), except that EPSCs were recorded from cultured hippocampal neurons (n = 11 cells). (E) Image showing a DRG neuron infected by ChR2-mCherry AAV2/9 virus and a co-cultured DH neuron. (F) Upper, diagram showing the protocol for infection of DRG neurons by ChR2-mCherry virus and co-culture with DH neurons; lower, cartoon of EPSC recording from DH neurons co-cultured with ChR2-expressing DRG neurons. (G) Left, typical EPSCs from DH neurons induced by light stimulation (475 nm, 5 ms, at arrows) of co-cultured ChR2-expressing DRG neurons in Ca^2+^-free or 2 mM Ca^2+^-containing solution before (black) and after (red) exposure to 50 μM AP5 and 10 μM CNQX. The EPSCs of 0Ca^2+^ and 2Ca^2+^ were recorded in 2 different DH neurons; right, statistics of EPSC amplitude (n = 5 cells for 0Ca^2+^ and 14 for 2Ca^2+^). Data are shown as the mean ± s.e.m.; Wilcoxon test for B; paired Student’s *t* test for D and G; **p <0.01, ***p <0.001. Scale bars, 20 μm.

We next used an optogenetic approach to specifically activate DRG neurons by expressing channelrhodopsin-2 (ChR2) in DRG neurons with AAV2/9 virus before the co-culture with DH neurons (***Figure 4E-F***). The transient application of 475-nm light evoked action potentials in ChR2-expressing DRG neurons (***Figure 4-figure supplement 1***). Importantly, the light-activation of DRG neurons also induced EPSCs in DH neurons bathed in either 0Ca or 2Ca solution, and these were fully abolished by the blockade of ionic glutamate receptors with D-AP5 and CNQX (***Figure 4G***). Together, these findings demonstrate that an action potential *per se* is able to directly trigger presynaptic glutamate release independent of Ca^2+^ influx during the synaptic transmission from presynaptic DRG terminals to postsynaptic central DH neurons.

### CiVDS-EPSC is mediated by SNARE complex and N-type Ca^2+^-channels

To determine whether the CiVDS-EPSC is mediated by SNARE-dependent vesicular exocytosis, we performed DRG neuron-restricted knockout of synaptobrevin 2 (Syb2/VAMP2) by infecting DRG neurons from homozygous floxed Syb2-null mice with Cre recombinase-carrying AAV2/5 virus (***Figure 5A***). Western blots showed the complete loss of Syb2 in DRG neurons infected with Cre virus (***Figure 5B***). Strikingly, the CiVDS-EPSC in DRG–DH transmission was substantially reduced by the Syb2-knockout (***Figure 5C***). Thus, similar to CiVDS in somatic secretion (Chai et al., 2017), CiVDS-mediated presynaptic glutamate release occurs through SNARE-dependent vesicular exocytosis.

**Figure 5.**
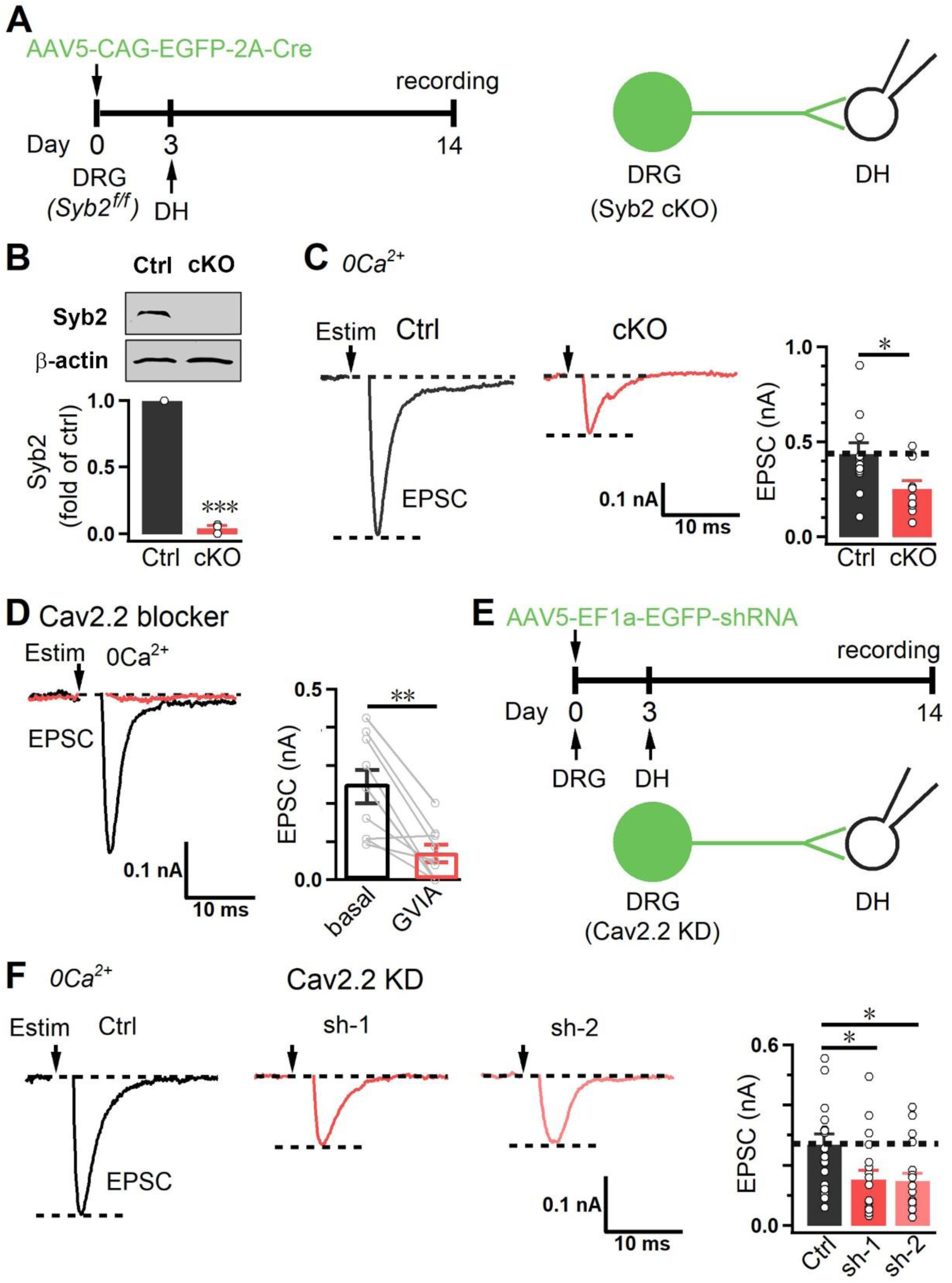
CiVDS-EPSCs are mediated by the SNARE complex and N-type Ca^2+^-channels. (A) Left, diagram showing the protocol for EPSC recording from DH neurons co-cultured with Syb2-knockout (cKO) DRG neurons; right, cartoon of EPSC recording from DH neurons co-cultured with Syb2-cKO DRG neurons. (B) Representative western blots (upper) and analysis (lower) of the expression levels of Syb2 in control (Ctrl) and Syb2-cKO DRG neurons (n = 3 per group). (C) Evoked EPSCs and statistics from DH neurons co-cultured with control (Ctrl) or Syb2-cKO DRG cells (cKO) in 0Ca^2+^ solution (n = 13 cells for Ctrl and 11 for cKO). (D) Evoked EPSCs and statistics from DH neurons co-cultured with DRG neurons before (black) and after (red) applying 1 μM ω-conotoxin-GVIA (GVIA) in 0Ca^2+^ bath (n = 9 cells). (E) Upper, diagram showing the protocol for EPSC recordings from DH neurons co-cultured with Cav2.2 knockdown (KD) DRG neurons; lower, cartoon of EPSC recording from DH neurons co-cultured with Cav2.2-KD DRG neurons. (F) Left panels, evoked EPSCs from DH neurons co-cultured with control (Ctrl) or Cav2.2-KD (sh-1/sh-2) DRG neurons in 0Ca^2+^ solution. Right panel, quantification of evoked EPSCs (n = 17 cells for Ctrl, 12 for sh-1, and 19 for sh-2). EPSCs were evoked by local electrical stimulation (Estim, at arrows). Data are shown as the mean ± s.e.m.; paired Student’s *t* test for B and D, unpaired Student’s *t* test for C, and Kruskal-Wallis test followed by Dunn’s multiple comparisons test for F; *p <0.05, **p <0.01, ***p <0.001.

We have shown that the N-type Ca^2+^ channel Cav2.2 serves as a voltage sensor for CiVDS in the somata of DRG and superior cervical ganglion neurons, while the L-type Ca^2+^ channel mediates CiVDS in adrenal slice chromaffin cells (Chai et al., 2017; Huang et al., 2019). To determine the voltage sensor for CiVDS in synaptic transmission of DRG-DH neurons, we blocked Cav2.2 with ω-conotoxin-GVIA (GVIA, 1 μM), which directly blocks the voltage-sensing ability of Cav2.2 (Chai et al., 2017; Ellinor, Zhang, Horne, & Tsien, 1994; Yarotskyy & Elmslie, 2009, 2010), and found that both CiVDS- and CDS-mediated EPSCs were remarkably decreased (***Figures 5D, Figure 5-figure supplement 1***). On the contrary, CiVDS-mediated EPSCs were insensitive to Cd^2+^, a well-known pore blocker of VGCCs (Chow, 1991; Huang et al., 2019; Tang et al., 2014) (***Figure 5-figure supplement 2***), confirming that the gating charge but not pore permeability is crucial for CiVDS-EPSCs. To further confirm the essential role of Cav2.2 in CiVDS-EPSCs, we adopted an shRNA-based knockdown (KD) approach. As validated previously (Chai et al., 2017), two independent Cav2.2-targeting shRNAs (sh-1 and sh-2) were delivered into DRG neurons by the AAV2/5 virus before co-culture with DH neurons (***Figure 5E***). Both the CiVDS- and CDS-mediated EPSCs in DH neurons were substantially decreased by Cav2.2-KD in presynaptic DRG neurons compared with those in control group (***Figure 5F, Figure 5-figure supplement 3***). Together, these findings indicate that Cav2.2 serves as a voltage sensor for CiVDS-mediated glutamate release during the synaptic transmission from DRG to DH neurons.

## Discussion

The classic view of neurotransmission is that impulse-induced presynaptic release (both somatic and terminal) occurs exclusively *via* CDS (Felmy et al., 2003; Geppert et al., 1994; Jackson & Chapman, 2006; Katz & Miledi, 1967; Meinrenken et al., 2003; Neher & Zucker, 1993; Sudhof, 2012). In somatic secretion, the CDS-only concept has recently been revised by a series of studies on CiVDS in peripheral sensory and sympathetic neurons (Huang et al., 2019; Liu et al., 2011; Moya-Diaz et al., 2019; Sforna et al., 2019; C. Zhang et al., 2004; C. Zhang & Zhou, 2002; Zheng et al., 2009), including its functions and mechanisms (Chai et al., 2017). In the present study, we show that CiVDS contributes to synaptic transmission in the central nervous system: at synapses between presynaptic DRG terminals and postsynaptic central DH neurons. The CiVDS-mediated EPSCs have a faster recycling rate and less short-term depression than CDS-mediated EPSCs. Similar to somatic CiVDS, the machinery governing CiVDS-mediated synaptic transmission includes Cav2.2 (voltage sensor) and Syb2-SNARE (fusion pore) (***Figure 6***).

**Figure 6.**
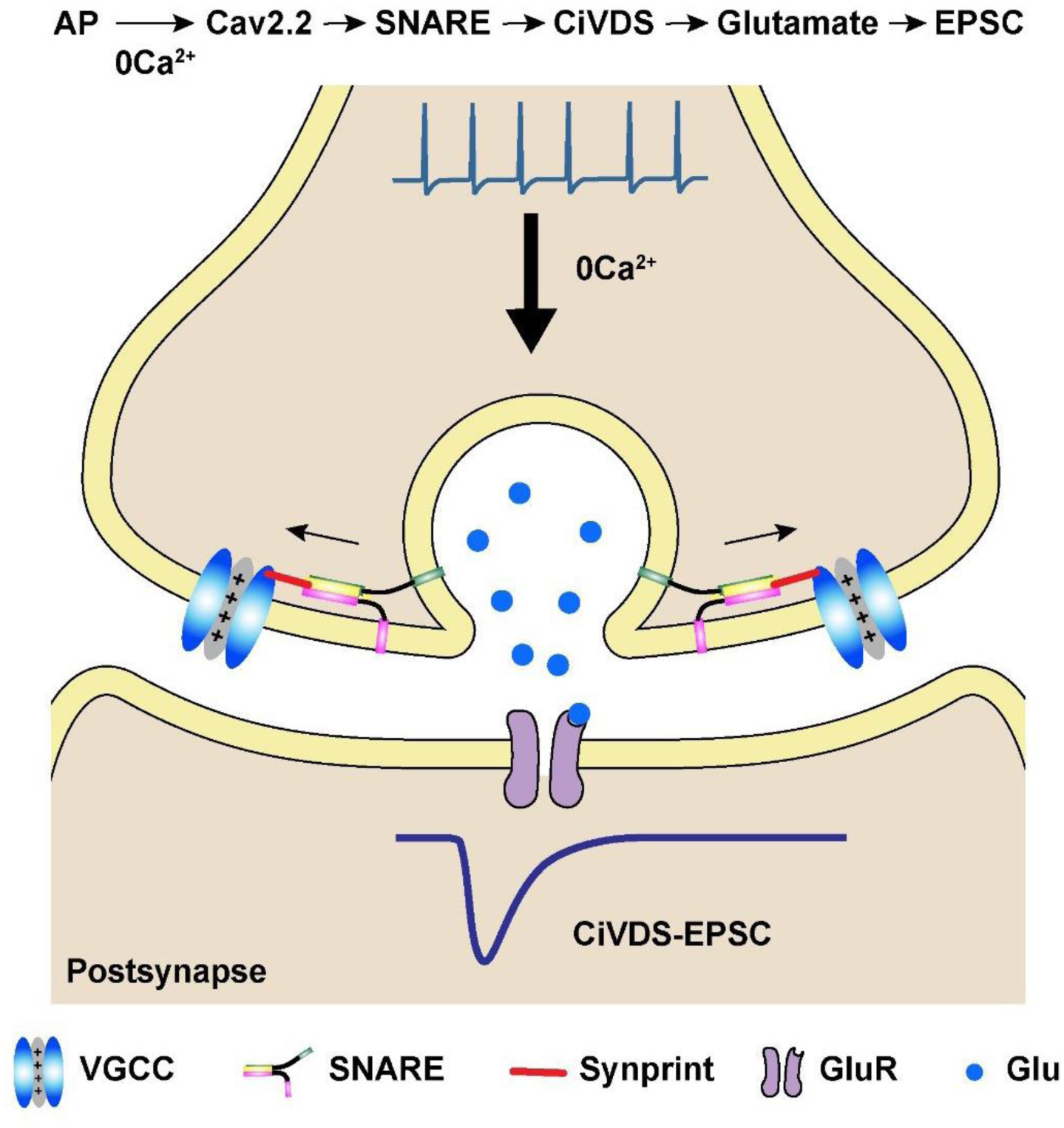
A model of the CiVDS-mediated EPSC in synaptic transmission. In Ca^2+^-free bath, an action potential activates the voltage-gated Ca^2+^ channel (VGCC/Cav2.2) and triggers presynaptic glutamate release through Ca^2+^-independent but voltage-dependent secretion (CiVDS), and activates an excitatory postsynaptic current (CiVDS-EPSC). In physiological solution containing Ca^2+^, however, both CiVDS- and Ca^2+^-dependent secretion (CDS)-mediated glutamate release contribute to a larger evoked postsynaptic EPSC (not shown).

The major finding of the present work is that CiVDS contributes to the central synaptic transmission between co-cultured presynaptic DRG and postsynaptic DH neurons. This is supported by the following evidence: (1) EPSCs were recorded in DRG-DH (***Figure 1A,C***) but not hippocampal neurons (***Figure 1B***) in 0Ca solution; (2) field stimulation triggered a [Ca^2+^]_i_ rise only in Ca^2+^-containing solution (***Figures 1A, Figure 1-figure supplement 1***); (3) in 2Ca solution, BAPTA reduced the EPSC amplitude to that in 0Ca (***Figure 1C-D***); (4) the evoked CiVDS-EPSCs were abolished by antagonists against AMPA receptors (CNQX) and NMDA receptors (AP5) (Trussell et al., 1993; C. Wang et al., 2016) (***Figure 3A-B***); (5) the decay of evoked CiVDS-EPSCs was greatly slowed by the AMPA receptor enhancer CTZ (Fucile et al., 2006) (***Figure 3C-D***); (6) Spy-pHluorin live-imaging confirmed CiVDS from the presynaptic terminals of DRG neurons but not hippocampal neurons (***Figure 2, Figure 2-video 1-4***); (7) paired patch-clamp recordings revealed that EPSCs between DRG-DH neurons were independent of extracellular and intracellular Ca^2+^, while the EPSCs in hippocampal neurons were completely blocked in the 0Ca condition (***Figure 4A-D***); (8) optogenetic stimulation of DRG neurons triggered EPSCs in either 0Ca or 2Ca solution, confirming that presynaptic glutamate release from DRG neurons is responsible for both the CiVDS-EPSCs and CDS-EPSCs recorded in DH neurons (***Figure 4E–G***); (9) SNARE is the CiVDS fusion machinery of the EPSCs, which were attenuated by knockout of the SNARE protein Syb2 in DRG neurons (***Figure 5A–C***); and (10) Cav2.2 is the voltage-sensor of CiVDS-EPSCs, as they were attenuated by either RNAi knockdown or an antagonist GVIA, which inhibits the voltage-sensor of Cav2.2 (Chai et al., 2017; Ellinor et al., 1994; Yarotskyy & Elmslie, 2009, 2010), but were insensitive to open Ca^2+^ channel blocker Cd^2+^ (Chow, 1991; Huang et al., 2019; Tang et al., 2014) (***Figures 5D–F, Figure 5-figure supplement 2***). Together, CiVDS contributes to the EPSCs between DRG and DH neurons in physiological solution and in response to physiological stimulation.

The second finding is that both CiVDS and CDS make essential and complementary contributions to the total EPSC (CDS + CiVDS) of DRG–DH synaptic transmission under physiological conditions: 2 mM Ca^2+^ bath solution and ≤20 Hz stimulation (Fang et al., 2005; Xu et al., 2000; Zheng et al., 2009). This is supported by the following evidence: (1) compared to the CDS-EPSC, the CiVDS-EPSC had a faster vesicle recycling rate (***Figure 1E-F***) and less short-term depression (***Figure 1G-H***); (2) CDS [EPSC(2Ca) – EPSC(0Ca)] and CiVDS [EPSC(0Ca)] contributed similarly (∼50% each) to the total EPSC during synaptic transmission following single-pulse/low-frequency stimulation (≤0.2 Hz) (***Figure 1D,F***); (3) CiVDS became dominant (∼87%) in the total EPSC following “painful” high-frequency stimulation (10-20 Hz) (Fang et al., 2005; Xu et al., 2000; Zheng et al., 2009), as CDS was attenuated more than CiVDS in the total EPSC (***Figure 1G-I***) due to much slower vesicle recycling of the CDS-EPSC (Neher & Sakaba, 2008; L. Y. Wang, Neher, & Taschenberger, 2008; B. Zhang et al., 2011); (4) single-pulses or low frequency (≤0.2 Hz) triggered both CDS and CiVDS in synaptic release (***Figure 1***), while only CiVDS (C. Zhang & Zhou, 2002; Zheng et al., 2009) but not CDS (Huang et al., 2019; Liu et al., 2011; Zheng et al., 2009) was triggered during somatic secretion; and (5) a “painful” burst of pulses (10-20 Hz) (Fang et al., 2005; Xu et al., 2000; Zheng et al., 2009) triggered both CiVDS and CDS in synaptic transmission (***Figure 1***), as well as somatic exocytosis (C. Zhang & Zhou, 2002; Zheng et al., 2009). Together, CiVDS and CDS make essential and complementary contributions to both synaptic EPSCs (***Figure 1***) and somatic transmitter release (Chai et al., 2017) in the DRG-DH sensory system.

The present study reported CiVDS-mediated synaptic transmission in a central synapse between DRG and DH neurons. In contrast, release from the presynaptic terminal of the calyx of Held is abolished after blocking Ca^2+^ influx (Felmy et al., 2003). Future work is needed to determine: (1) whether CiVDS-EPSCs occur in the synaptic transmission of other neural circuits; (2) whether CiVDS-IPSCs exist; (3) whether CiVDS-EPSCs occur in brain slices and *in vivo*; and (4) the physiological relevance of CiVDS-EPSCs.

In summary, the present work extends the occurrence of CiVDS from peripheral neuronal somata and neuroendocrine cells to a central presynaptic terminal, and demonstrates CiVDS-EPSCs in a rat/mouse model of cultured DRG–DH neurons, implying potential physiological roles of CiVDS in synaptic transmission in other mammalian neural circuits.

## Materials and methods

### Animals

The use and care of animals was approved and directed by the Institutional Animal Care and Use Committee of Peking University and the Association for Assessment and Accreditation of Laboratory Animal Care. We used CRISPR/Cas9-mediated one-step genomic engineering to generate Synaptobrevin-2 floxed mice (Yang et al., 2013). A mixture of guide RNAs (CTCTGGTGATAGGCGGATCCAGG, AGGGTTCCTAGACGAACACCAGG), the corresponding donor DNA and the Cas9 protein was micro-injected into the fertilized eggs, followed by embryonic transplantation. Two LoxP sites were inserted into the 5’ intron of exon 3 and 3’ intron of exon 4 in mouse synaptobrevin-2 gene donor fragment (Gene ID: 22318). All animals were housed in an animal facility under a 12-h light/dark cycle at 22 ± 2°C with food and water available *ad libitum*. Sprague-Dawley rats were used for all experiments except that floxed Syb2-null mice were used for Syb2-KO experiments.

### Cell culture

Three to four-day old postnatal Sprague-Dawley rats or floxed Syb2-null mice were used for the culture of dorsal root ganglion (DRG) neurons and 15–16 day embryos of Sprague-Dawley rats were used for the culture of dorsal horn (DH) neurons. Postnatal day 0 Sprague-Dawley rats were used for the culture of hippocampal neurons.

DRG isolation and neuronal culture were performed as previously described (Chai et al., 2017). DRG ganglia were dissected from the spine and placed in cold L15 medium (Gibco). After removing attached tissue, the ganglia were cut into several pieces and incubated in Dulbecco’s modified Eagle’s medium (DMEM)/F12 (Gibco) containing trypsin (0.2-0.3 mg/ml) and collagenase (1 mg/ml) for 40 min at 37°C under 5% CO_2_. After that, the pieces were washed twice with 2 ml DMEM/F12 and dissociated into single cells by 8–10 bouts of trituration. For confocal imaging, cells were collected and transfected with Synaptophysin (Spy)-pHluorin plasmid (a kind gift from Dr. G. Miesenböck, University of Oxford) by using the NeonTM (100-μl) electroporation system (MPK10096, Invitrogen). Then the cell suspension was placed on 0.1% poly-L-lysine pretreated coverslips and maintained in Neurobasal supplemented with 2% B27 and 0.5 mM L-glutamine (all from Gibco).

DH neurons were isolated and cultured as previously described (Zheng et al., 2009). The spinal cord was dissected from 15–16-day embryos and placed in cold D-Hanks solution. After removing attached tissue, the dorsal spinal cord was incubated in 0.25% trypsin solution for 15 min at 37°C under 5% CO_2_. After that, the dorsal cord was washed twice with 2 ml Neurobasal supplemented with 2% B27 and 0.5 mM L-glutamine (all from Gibco) and dissociated into single cells by 8–10 bouts of trituration. To co-culture DRG and DH neurons, the suspension of DH neurons was evenly placed onto pre-cultured DRG neurons. All cells were maintained in Neurobasal supplemented with 2% B27, 0.5 mM L-glutamine (all from Gibco), 10 ng/ml nerve growth factor (NGF), 50 ng/ml brain-derived neurotrophic factor (BDNF), 50 ng/ml glial cell line-derived neurotrophic factor (GDNF), and 5 μM cytosine arabinoside. The mixture of NGF, BDNF, and GDNF helped to maintain CiVDS for >2 weeks. The neurons were used between days 10-12 after the start of co-culture.

For hippocampal cultures, hippocampal neurons were prepared as previously described (C. Wang et al., 2018). Briefly, hippocampi were dissected from neonatal rats and placed in cold D-Hanks solution. After removing attached tissue, the hippocampi were digested in 0.25 % trypsin for 15 min at 37°C under 5% CO_2_. After that, the tissue was washed twice with 2 ml Neurobasal supplemented with 2% B27 and 0.5 mM L-glutamine (all from Gibco), and dissociated by 8–10 bouts of trituration. Then the cells were placed on 0.1% poly-L-lysine-coated coverslips and maintained in DMEM (Gibco) supplemented with 10% FBS for 3 h, which was then replaced by Neurobasal supplemented with 2% B27 and 0.5 mM L-glutamine. Experiments were performed between days 14-16 after the start of culture. For confocal imaging, cultures at day 5 were transfected with Spy-pHluorin plasmid by using a calcium-phosphate transfection method. Briefly, plasmids in a 250 mM CaCl_2_ solution were slowly added to Hank’s balanced salt solution (HBSS) and incubated at room temperature for 25 min. The mixture was then added to the culture and incubated for 15 min. The cells were washed with MgCl_2_-containing medium and then maintained in the original medium. Confocal imaging was performed at day 13-14.

### Virus infection

For virus infection, the virus was added to the culture solution at the beginning of DRG neuronal culture. After 3 days, the virus-containing culture medium was fully replaced and then the DH neurons were co-cultured with the DRG neurons. The adeno-associated virus vector carrying CAG-hChR2(H134R)-mCherry (AAV2/9) was used for optogenetics and CAG-EGFP-2A-Cre (AAV2/5) for Syb2-knockout. For Cav2.2 knockdown, the nucleotide target sequences GG ACA TTT CTG CAA GCC TTA A (shCav2.2-1) and GC TAC TTC CGG TCT TCC TTC A (shCav2.2-2) were integrated into adeno-associated virus (AAV2/5) to silence the expression of Cav2.2. All the viruses were from Shanghai Heyuan Biotech (China).

### Electrophysiology

Excitatory postsynaptic currents (EPSCs) were recorded under the whole-cell configuration using an EPC10/2 amplifier controlled by Patchmaster software (HEKA Elektronik) as described previously (B. Zhang et al., 2011; Zheng et al., 2009). The membrane potential was clamped at –70 mV, and pipette resistance was controlled to ∼10 MΩ for EPSC recordings. The standard external bath solution contained (in mM): 150 NaCl, 5 KCl, 2.5 CaCl_2_, 1 MgCl_2_, 10 H-HEPES, and 10 D-glucose, pH 7.4. The Ca^2+^-free solution for CiVDS recording was the same, except that 2.5 mM Ca^2+^ was replaced by 1 mM EGTA. For EPSC recording, 100 μM picrotoxin was added to the bath to block IPSCs (inhibitory postsynaptic currents). The standard intracellular pipette solution contained (in mM): 145 K-gluconate, 5 KCl, 4 MgCl_2_-6H_2_O, 10 H-HEPES, 5 EGTA, and 2 QX314, pH 7.2. Standard intracellular solution with CsCl containing (in mM) 153 CsCl, 1 MgCl_2_, 10 H-HEPES, 4 Mg-ATP, 5 QX314, pH7.2 was used for a stable recording in Syb2-KO and optogenetic experiments. For EPSC recordings, local electrical field or light stimulation was applied. Field electrical stimulation (Estim) was used to evoke action potentials *via* a laboratory-made bipolar microelectrode (150 μm in diameter) connected to an electronic stimulator (Nihon Kohden, SEN-3201) (B. Zhang et al., 2011).

In paired patch-clamp recordings, the presynaptic DRG neuron was in whole-cell current-clamp mode and the postsynaptic dorsal horn neuron was in whole-cell voltage-clamp mode. For the current-clamp of DRG neurons, the intracellular solution contained (in mM) 135 K-gluconate (115 when adding 10 mM BAPTA), 5 KCl, 4 MgCl_2_, 3 MgATP, 0.3 Na_2_GTP, and 10 Na_2_-phosphocreatine, 10 HEPES, pH 7.2. For the voltage-clamp DH neuron, standard intracellular solution (but CsCl replacing KCl) was used. For light stimulation, 475-nm light was generated by a laser (VD-IIIA, Beijing Viasho Technology, China).

All recordings were made at room temperature (25°C). Igor software (Wavemetrics, Lake Oswego, OR) was used for all offline data analysis. Series conductance and membrane conductance were used to monitor the seal condition during patch-clamp recordings.

### Ca^2+^ imaging

Ca^2+^ imaging was conducted as described previously (Huang et al., 2019; Shang et al., 2016). Cytosolic Ca^2+^ was measured with the Ca^2+^ indicator Cal-520 (21130, AAT Bioquest). Co-cultured DRG and DH neurons were loaded with 0.5 μM Cal-520 AM for 30 min at 37°C and then washed 3 times with 2 mM Ca^2+^ solution at room temperature. Then the cells were electrically simulated and imaged on an inverted confocal microscope (LSM710, Zeiss). The fluorescent signal was captured by excitation with a 488-nm laser and emission at 500–560 nm. Fluorescence intensity values from the images were calculated and analyzed using ImageJ. The cutoff for selection (5 a.u.) was 1.5-fold that of the full-width at half-maximum of the Gaussian distribution of fluorescence noise.

### Confocal live-imaging

Time-lapse images were captured at 0.5 s interval through the inverted confocal microscope (LSM710, Zeiss) with a 63 x oil-immersion objective lens (Zeiss). The fluorescent signal was captured by excitation with a 488-nm laser and emission at 500–560 nm at room temperature bathed in Ca^2+^-free or standard external solution. Electrical field stimulation was applied via an electrical stimulator (Nihon Kohden, SEN-3201). Fluorescence changes at individual boutons were monitored over time and calculated as ΔF/F_0_. Images were acquired for more than 200 s in total. The first 30 s before stimulation was used to establish a baseline. Data were analyzed offline with ImageJ.

### Protein preparation and western blotting

Cells were washed with phosphate-balanced saline (PBS) and homogenized on ice with lysate buffer (RIPA; C1503, Beijing Applygen Technologies Inc.), 1 mM PMSF, and 2% proteinase inhibitor (539,134; Calbiochem). The homogenates were centrifuged at 13,000 rpm for 10 min at 4°C and the supernatants were collected and boiled in SDS-PAGE buffer. Proteins were electrophoresed and transferred to nitrocellulose filter membranes (Pall Life Sciences). The membranes were blocked for 1 h with PBS containing 0.1% Tween-20 (v/v) and 5% fat-free milk (w/v), then incubated with primary antibodies at 4°C overnight in PBST containing 2% bovine serum albumin. After washing with 0.1% Tween-20 containing PBS (PBST), they were then incubated with secondary antibodies at room temperature for 1 h. Blots were scanned with an Odyssey infrared imaging system (LI-COR Biosciences) and quantified with ImageJ (National Institutes of Health, Bethesda, MD). The primary antibodies used were anti-Syb2 (104202, SYSY) and β-actin (A5316, Sigma). The secondary antibodies were IRDye 800CW goat anti-rabbit IgG (LIC-926-32211) and IRDye 680CW goat anti-mouse IgG (LIC-926-32220), both from LI-COR Biosciences.

### Statistical analysis

All experiments were performed with controls side-by-side and in random order, and were replicated at least three times. All data were collected at least every 2 weeks within 3 months window to maintain the stability of a data set. Sample sizes are consistent with those reported in similar studies. Multiple recordings from one cell with the identical stimulus protocol were considered as technical replications, which were averaged to generate a single biological replication representing value/data from one cell. No samples or recordings that provided successful measurements were excluded from analysis. Data are shown as the mean ± s.e.m. All the data were tested for normality by Shapiro-Wilk test before the statistical analysis. If the data passed the normality test, two-tailed Student’s *t* test was applied for comparison between two unpaired groups and the paired Student’s *t* test was used for comparison between two matched groups. If the data did not pass normality, Mann-Whitney test was applied for comparison between two unpaired groups, Wilcoxon matched-pairs signed rank test (Wilcoxon test) was applied for two matched groups and the Kruskal-Wallis test followed by Dunn’s multiple comparisons test was used when multiple groups were compared with one variable. All tests were conducted using Prism V7.0 (GraphPad Software, Inc.) and SPSS 20.0 (Statistical Package for the Social Sciences). Significant differences were accepted at p <0.05.

## Acknowledgements

We thank Drs. Mengping Wei and Chen Zhang (Capital Medical University) for advices on EPSC recordings and virus transfection *in vivo*, Jianguo Gu (UA Birmingham), Kun Yang (Jiangsu University) and Luyang Wang (Toronto University) for helpful discussions, and Iain C. Bruce (Peking University) for reading the manuscript. This work was supported by National Natural Science Foundation of China (31930061, 21790394, 31761133016, 31821091, 31330024, 31171026, 31327901, 32171233, 31670843 and 21790390), the National Key Research and Development Program of China (2016YFA0500401), the Shaanxi Natural Science Funds for Distinguished Young Scholars of China (2019JC-07), the Innovation Capability Support Program of Shaanxi Province, China (2021TD-37), and the China Postdoctoral Science Foundation (2020M680211, 2021T140014).

## Additional information

### Funding

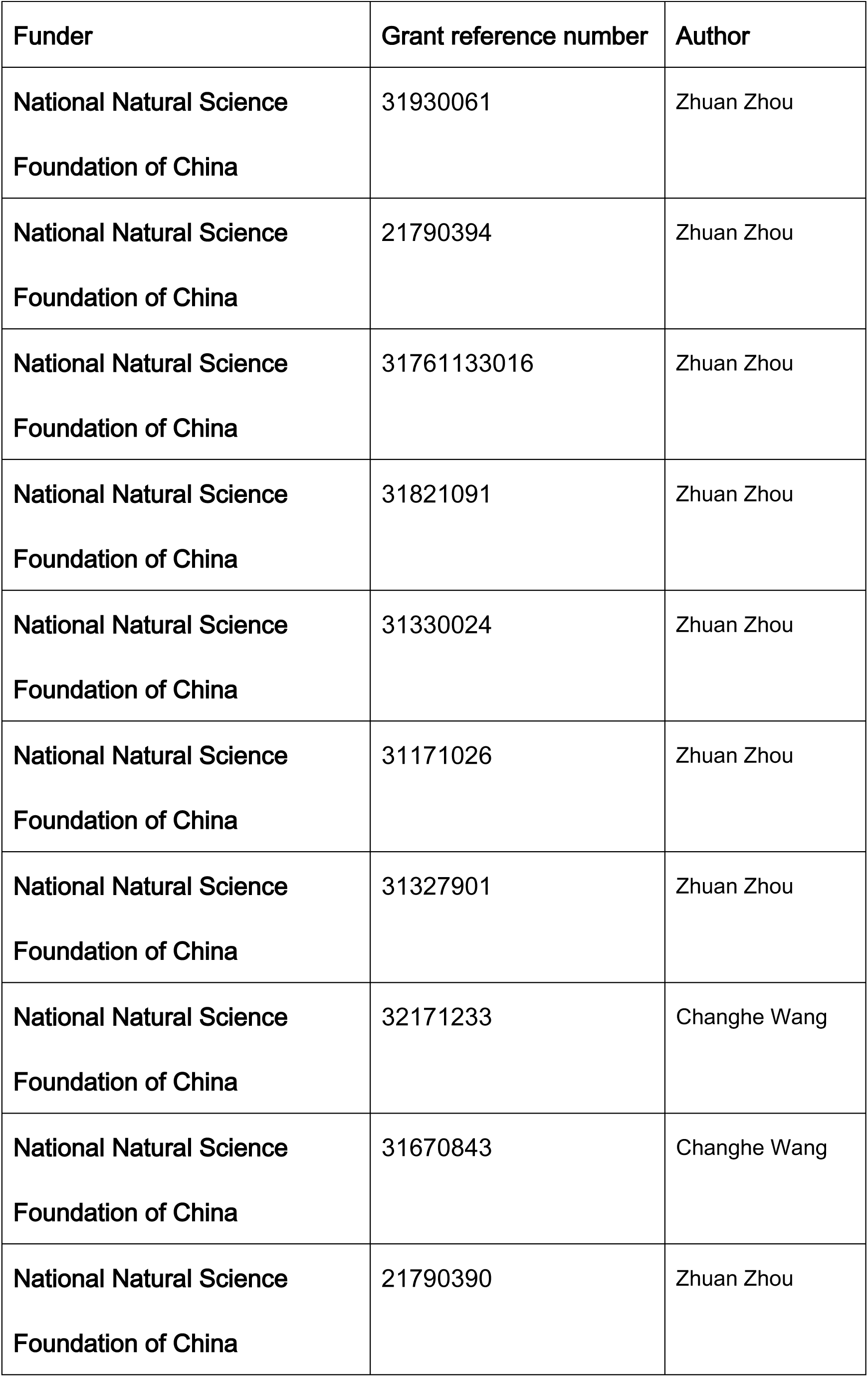
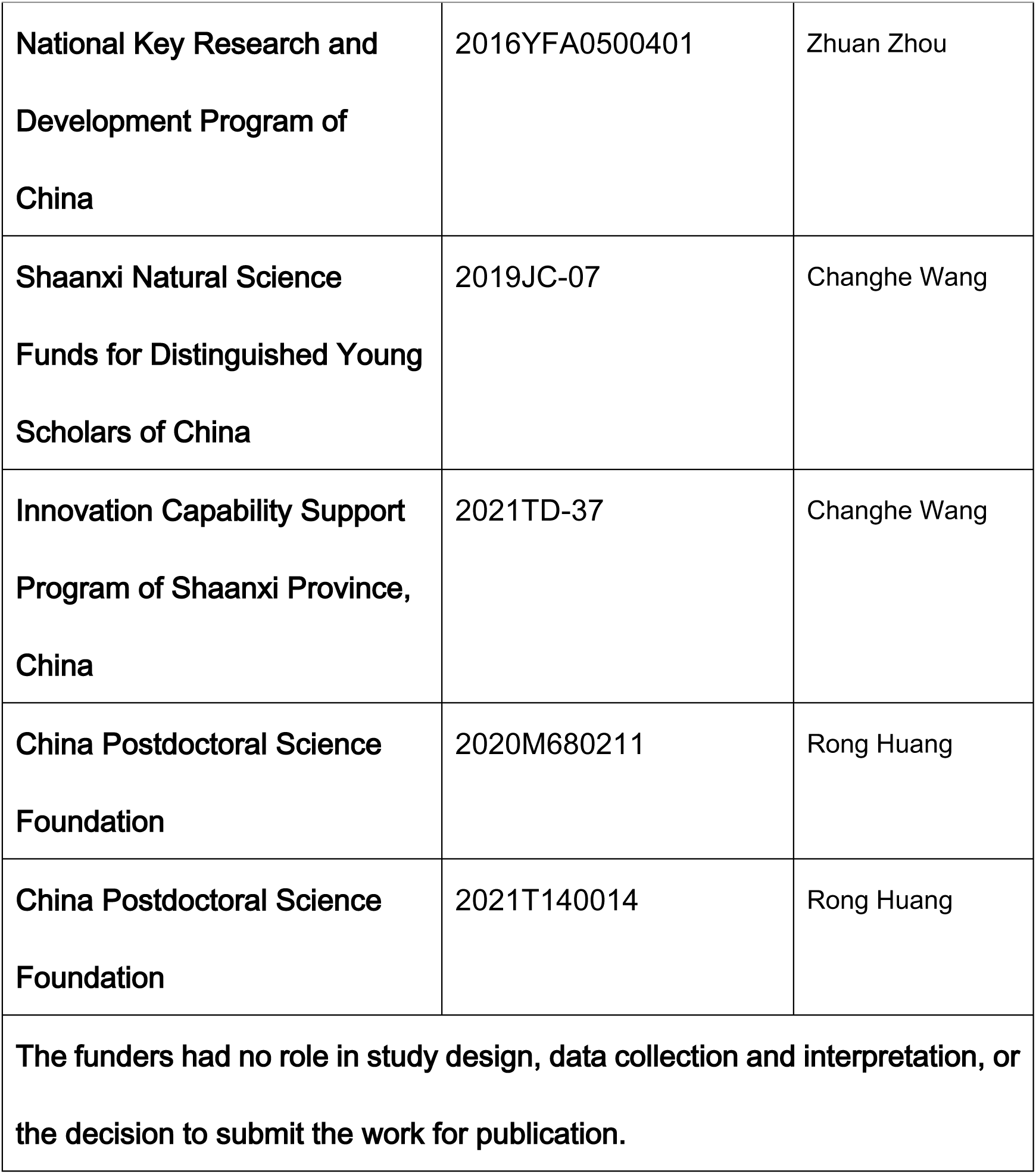

| **Funder** | **Grant reference number** | **Author** |
| --- | --- | --- |
| National Natural Science<br>Foundation of China | 31930061 | Zhuan Zhou |
| National Natural Science<br>Foundation of China | 21790394 | Zhuan Zhou |
| National Natural Science<br>Foundation of China | 31761133016 | Zhuan Zhou |
| National Natural Science<br>Foundation of China | 31821091 | Zhuan Zhou |
| National Natural Science<br>Foundation of China | 31330024 | Zhuan Zhou |
| National Natural Science<br>Foundation of China | 31171026 | Zhuan Zhou |
| National Natural Science<br>Foundation of China | 31327901 | Zhuan Zhou |
| National Natural Science<br>Foundation of China | 32171233 | Changhe Wang |
| National Natural Science<br>Foundation of China | 31670843 | Changhe Wang |
| National Natural Science<br>Foundation of China | 21790390 | Zhuan Zhou |
| National Key Research and Development Program of China | 2016YFA0500401 | Zhuan Zhou |
| Shaanxi Natural Science Funds for Distinguished Young Scholars of China | 2019JC-07 | Changhe Wang |
| Innovation Capability Support Program of Shaanxi Province, China | 2021TD-37 | Changhe Wang |
| China Postdoctoral Science Foundation | 2020M680211 | Rong Huang |
| China Postdoctoral Science Foundation | 2021T140014 | Rong Huang |
| The funders had no role in study design, data collection and interpretation, or the decision to submit the work for publication. |  |  |

### Author contributions

Y.W., R.H., X.D. and Y.H. performed and analyzed the experiments with the help of Y.X., J.L., X.J., X.W., Z.Q., Y.L., B.L. and F.Z.; Y.Z. and P.C. generated floxed Syb2-null mice; Z.Z., Z.C. and C.W. designed the work and wrote the manuscript with inputs from all authors. Z.Z. supervised the project.

---

**Figure 1—figure supplement 1.**
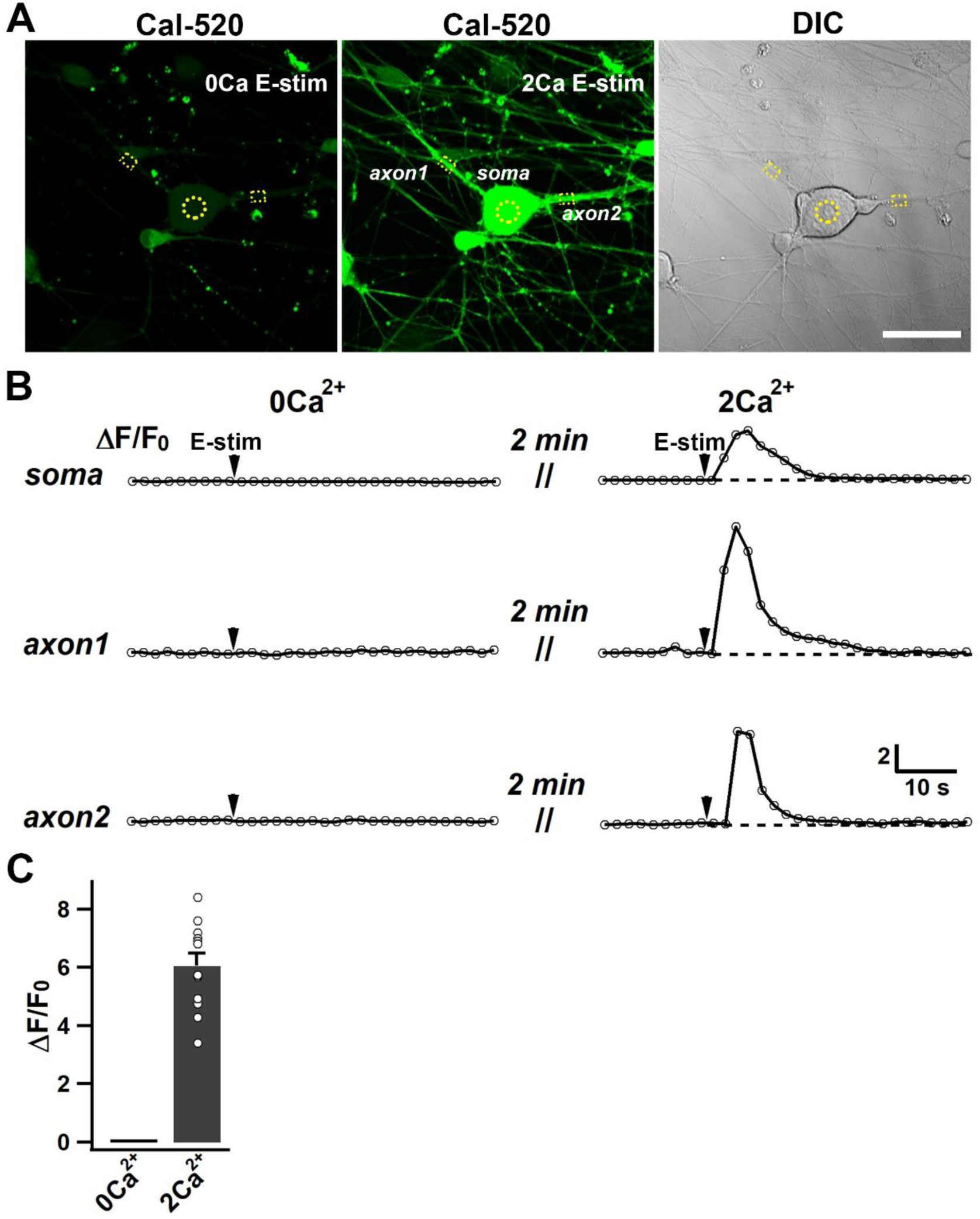
Absence of intracellular Ca^2+^ signal (ΔF/F_0_) in DRG neurons in Ca^2+^-free solution. (A) Representative images showing the loading of Cal-520 in co-cultured DRG and DH neurons (scale bar, 50 μm; DIC, differential interference contrast). (B) Ca^2+^ signal ΔF/F_0_ of Cal-520 fluorescence induced by electrical stimulation (E-stim at arrows) in Ca^2+^-free (left) and 2 mM Ca^2+^-containing solution (right) from typical regions labeled in (A). (C) Statistics of [Ca^2+^]_i_ rise as in (B) (n = 12 cells). No [Ca^2+^]_i_ rise in Ca^2+^-free solution was detectable. Data are shown as the mean ± s.e.m. Student’s *t* test for (C); ***p <0.001.

**Figure 1—figure supplement 2.**
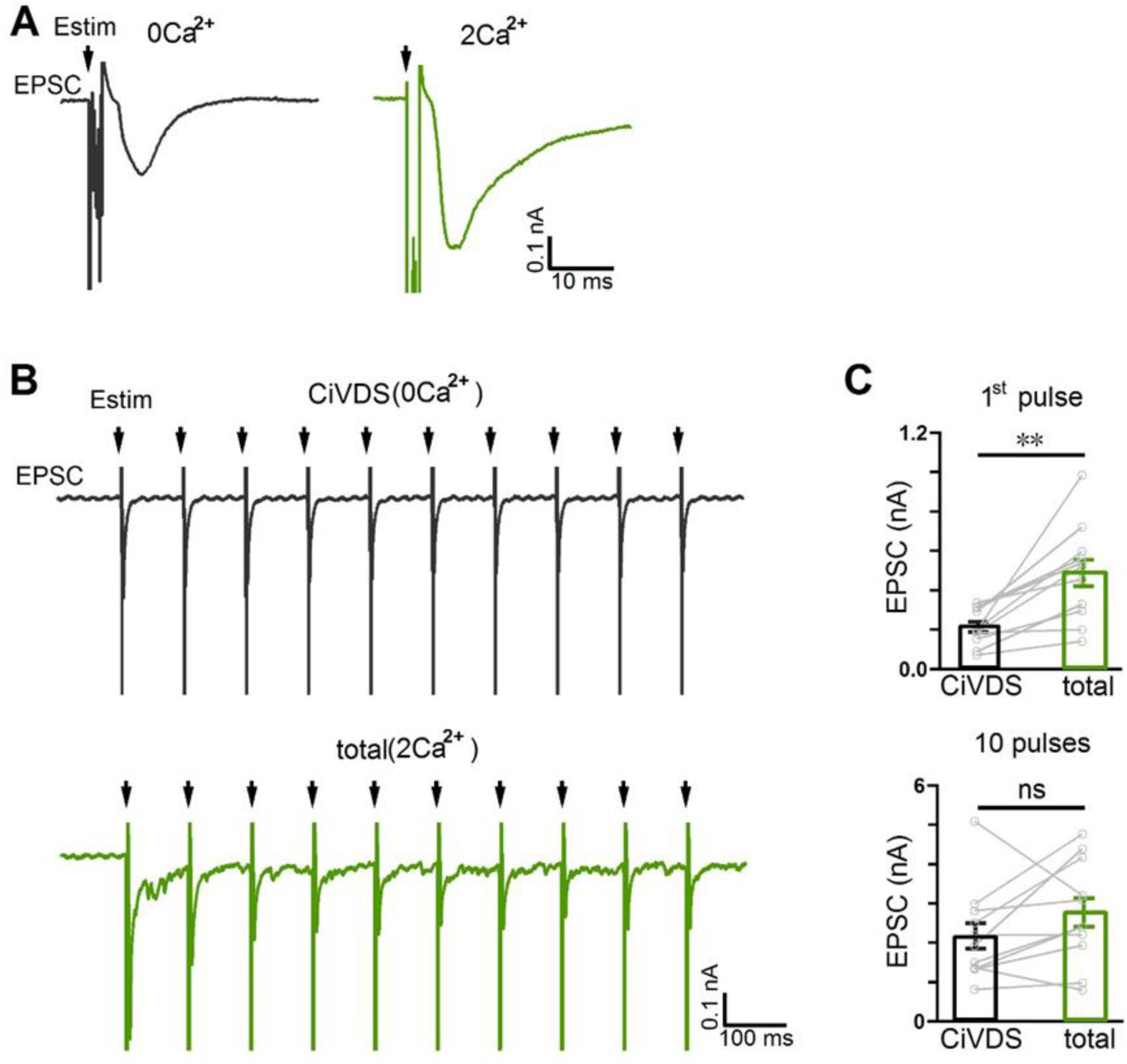
EPSCs evoked by local-field electrical stimuli from DH neurons co-cultured with DRG neurons. (A) Original recordings of EPSCs including artifacts in response to a single electrical stimulus (Estim, arrows) in 0Ca^2+^ (black, left) or 2Ca^2+^ solution (green, right). The EPSCs of 0Ca^2+^ and 2Ca^2+^ were recorded in different DH neurons. (B) As in (A) for CiVDS (0Ca^2+^) or total (CiVDS +CDS, 2Ca^2+^), but in response to a 10-Hz train of 10 pulses. The EPSCs of 0Ca^2+^ and 2Ca^2+^ were recorded in different DH neurons. (C) Statistics of CiVDS-EPSC (0Ca^2+^, black) and total (CDS + CiVDS)-EPSC (2Ca^2+^, green) evoked by single (first) pulse or 10 pulses during 10-Hz train stimulation as in (B) (n = 11 cells). Data are shown as mean ± s.e.m.; paired Student’s *t* test for (C), **p <0.01, ns, not significant.

**Figure 3—figure supplement 1.**
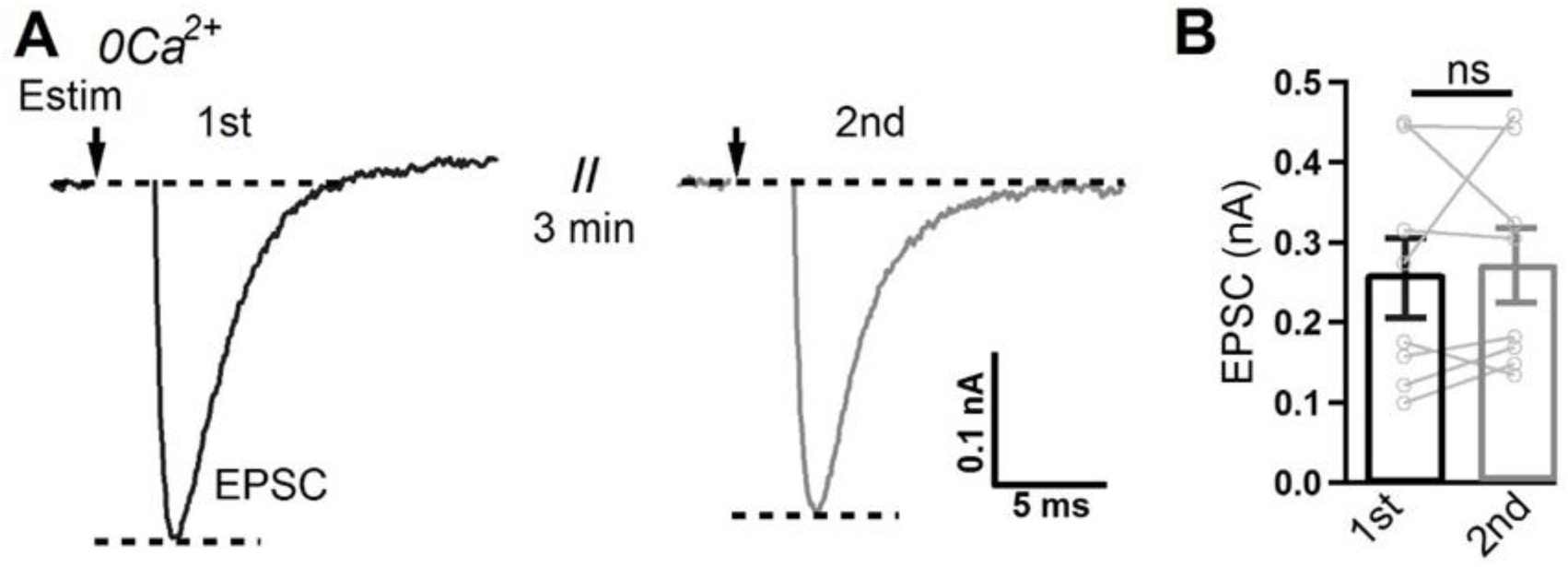
CiVDS-EPSC is reproducible during a repeated stimulus. (A) CiVDS-EPSCs induced by electrical stimulation (Estim, arrows) repeated after a 3-min interval in a DH neuron co-cultured with DRG neurons in 0Ca^2+^ solution. (B) Quantification of EPSC amplitude as in (A) (n = 8 cells). Data are shown as the mean ± s.e.m.; paired Student’s *t* test for (B); ns, not significant.

**Figure 4—figure supplement 1.**
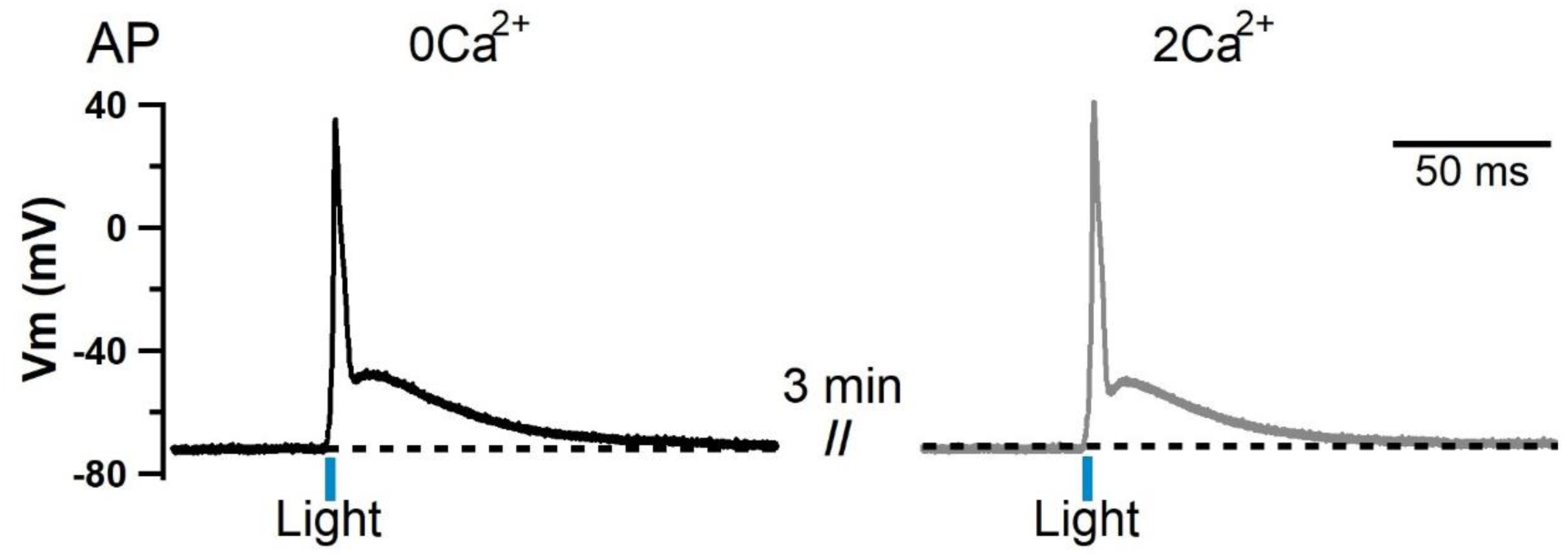
Light-evoked action potentials in DRG neurons expressing ChR2. Representative action potentials (APs) evoked by light stimulation (475 nm, 5 ms) in a ChR2-expressing DRG neuron in 0Ca^2+^ (left) and 2Ca^2+^ (right) solutions.

**Figure 5—figure supplement 1.**
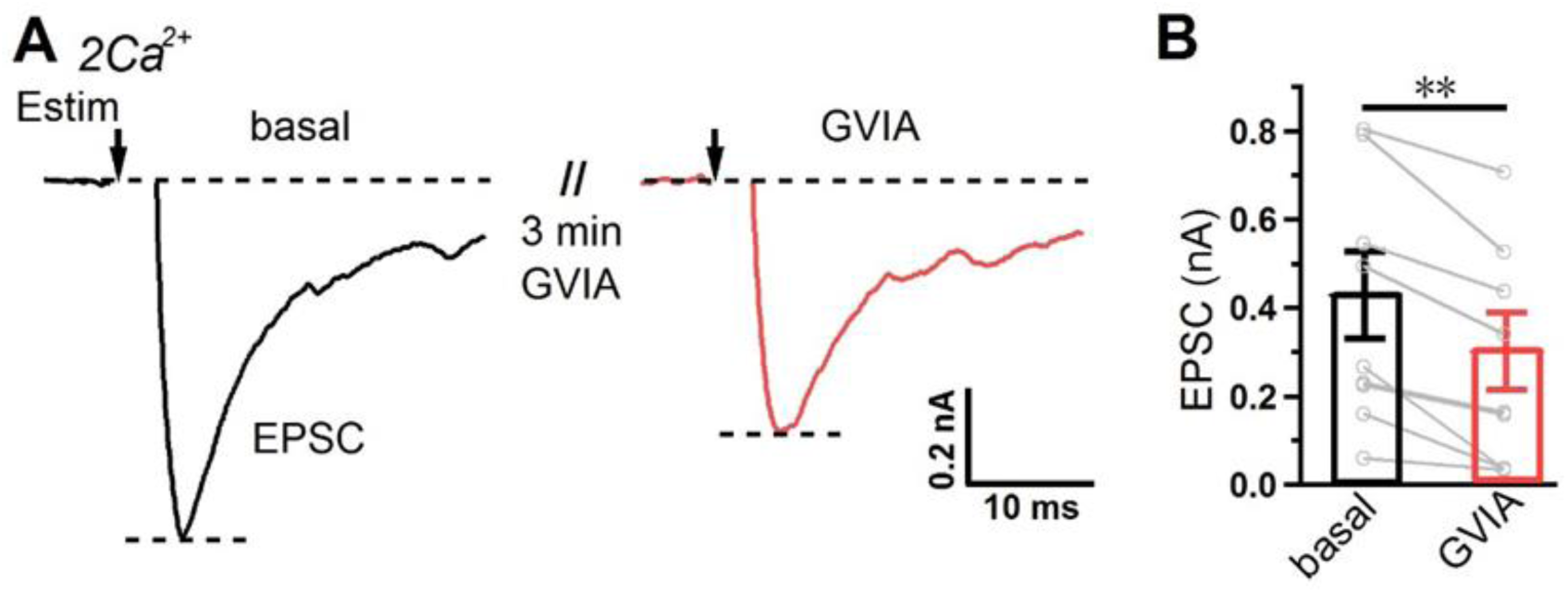
EPSCs in DH neurons co-cultured with DRG neurons are partially blocked by the Cav2.2 inhibitor GVIA. (A) EPSCs induced by electrical stimulation (Estim, at arrows) in a DH neuron co-cultured with DRG neurons before (black) and after (red) 1 μM ω-conotoxin-GVIA (GVIA) application in 2Ca^2+^ solution. (B) Quantification of EPSC amplitudes as in (A) (n = 9 cells). Data are shown as the mean ± s.e.m.; paired Student’s *t* test for (B); **p <0.01.

**Figure 5—figure supplement 2.**
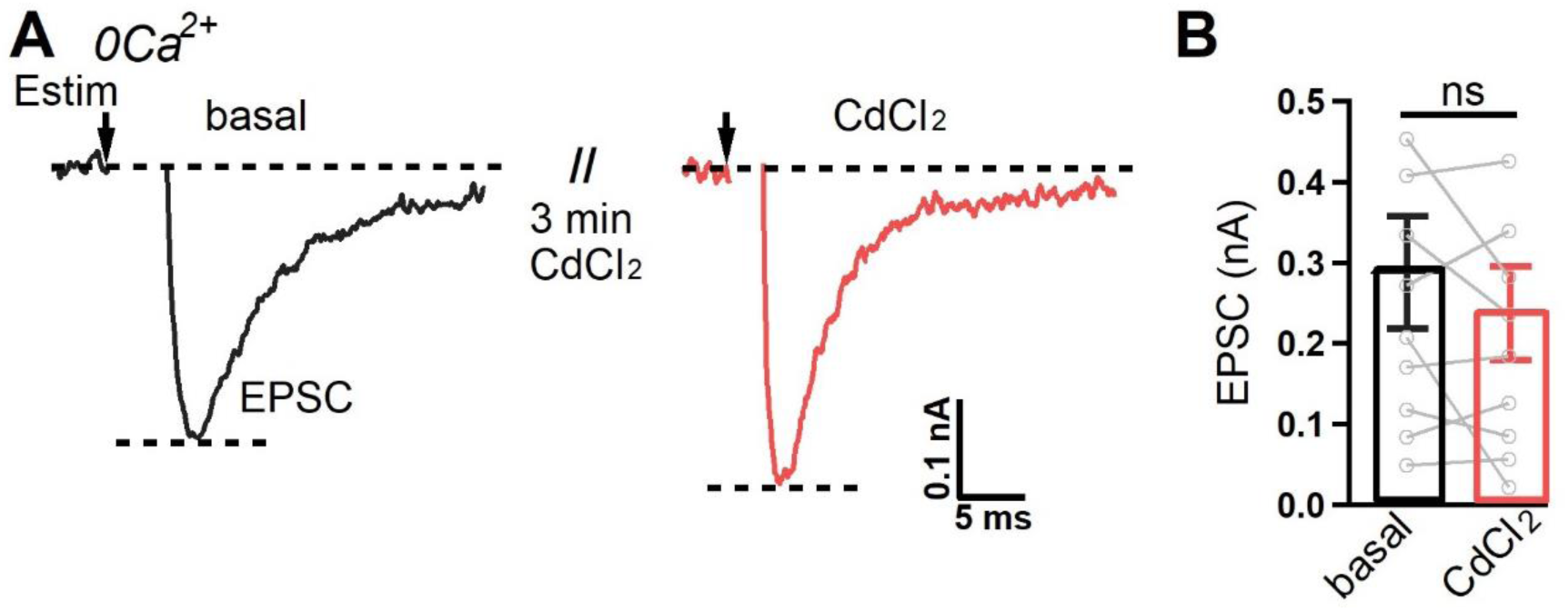
CiVDS-EPSCs in DH neurons co-cultured with DRG neurons are insensitive to CdCl_2_. (A) EPSCs induced by electrical stimulation (Estim, arrows) in a DH neuron co-cultured with DRG neurons before (basal) and after 100 μM CdCl_2_ application (CdCl_2_) in 0Ca^2+^ solution. (B) Quantification of EPSC amplitudes as in (A) (n = 10 cells). Data are shown as the mean ± s.e.m.; paired Student’s *t* test for (B); ns, not significant.

**Figure 5—figure supplement 3.**
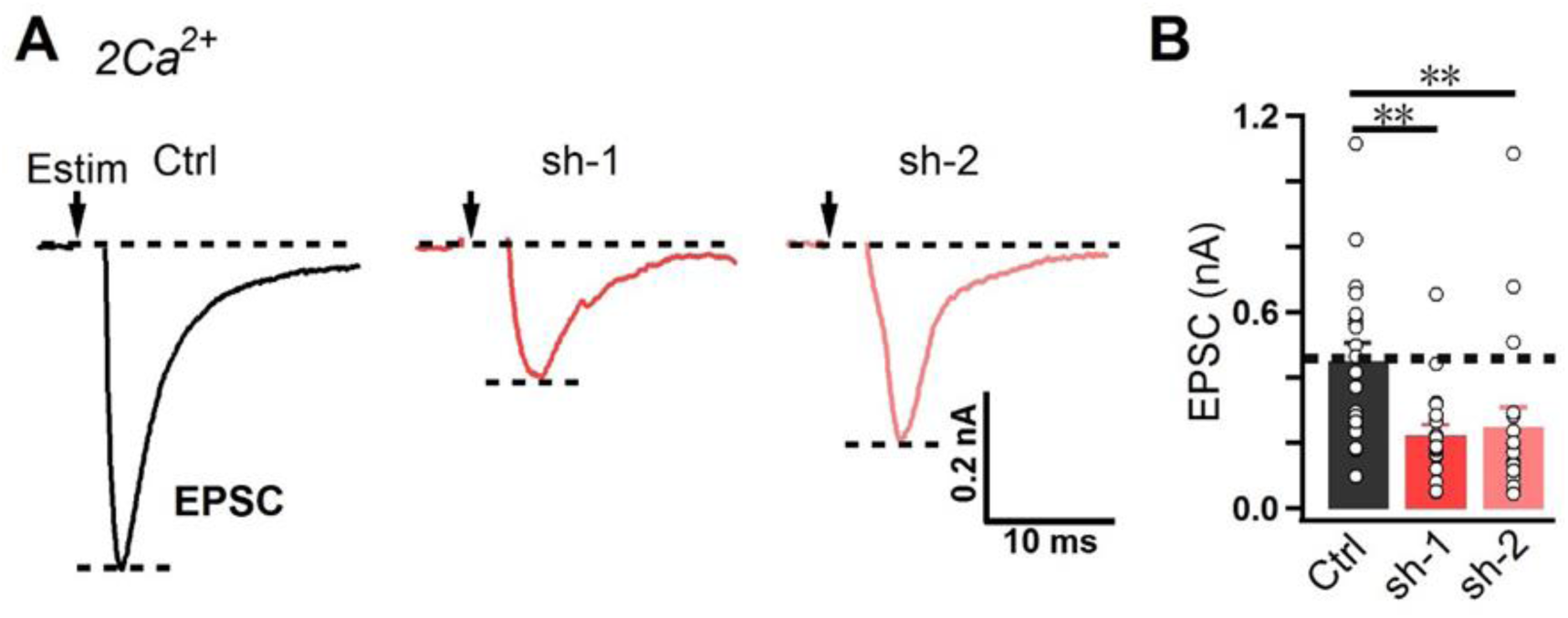
EPSCs in DH neurons co-cultured with DRG neurons are partially blocked by Cav2.2 knockdown. (A) Typical EPSCs induced by electrical stimulation (Estim, arrows) in DH neurons co-cultured with control (Ctrl) or Cav2.2-KD virus (sh-1/sh-2)-infected DRG neurons in 2 mM Ca^2+^-containing solution. (B) Quantification of EPSC amplitude as in (A) (n = 18 cells for Ctrl, 15 for sh-1, and 19 for sh-2). Data are shown as the mean ± s.e.m.; Kruskal-Wallis test followed by Dunn’s multiple comparisons test for (B); **p <0.01.

**Figure 2-video 1.**
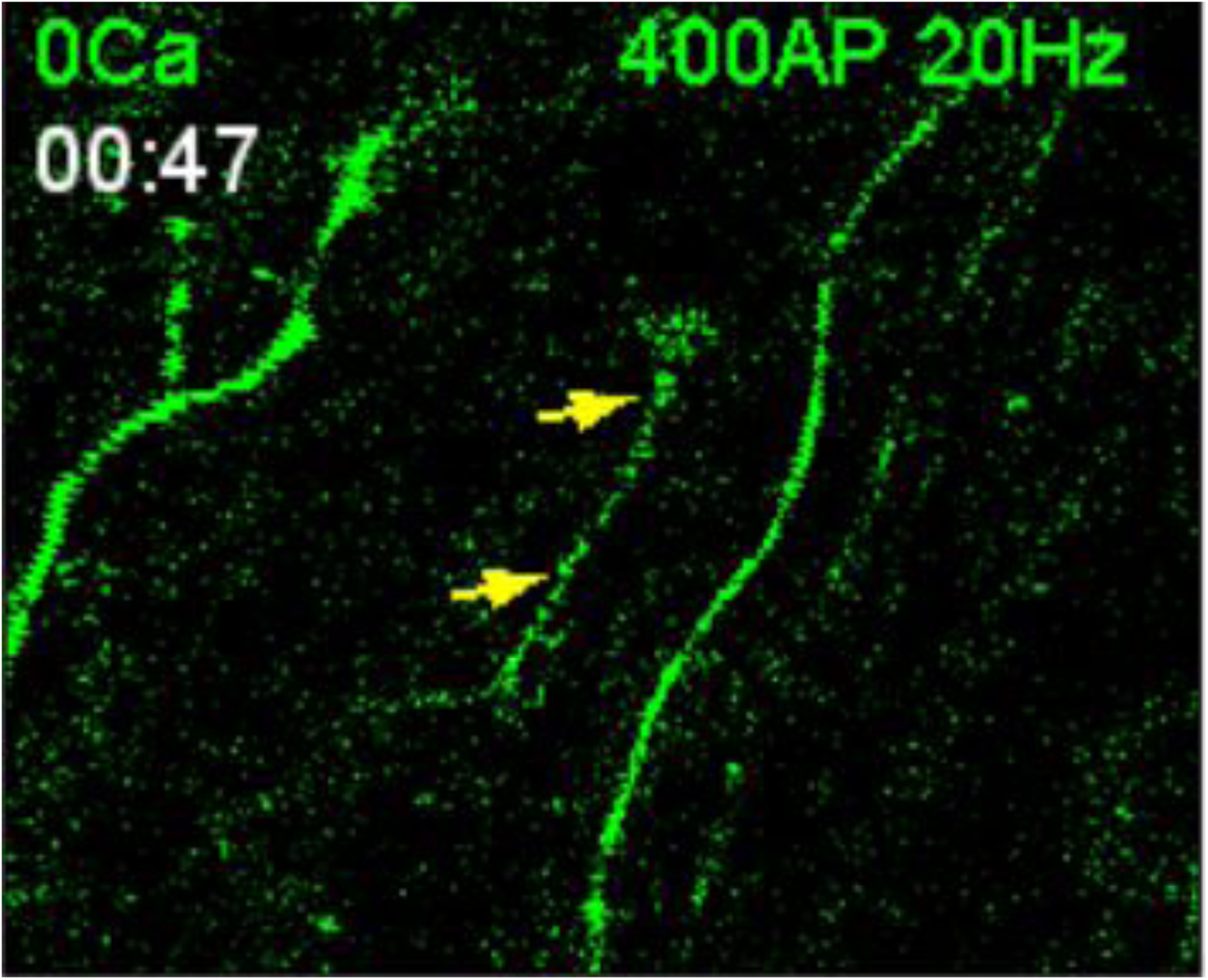
Spy-pHluorin signal from DRG-DH neuron in 0Ca solution.

**Figure 2-video 2.**
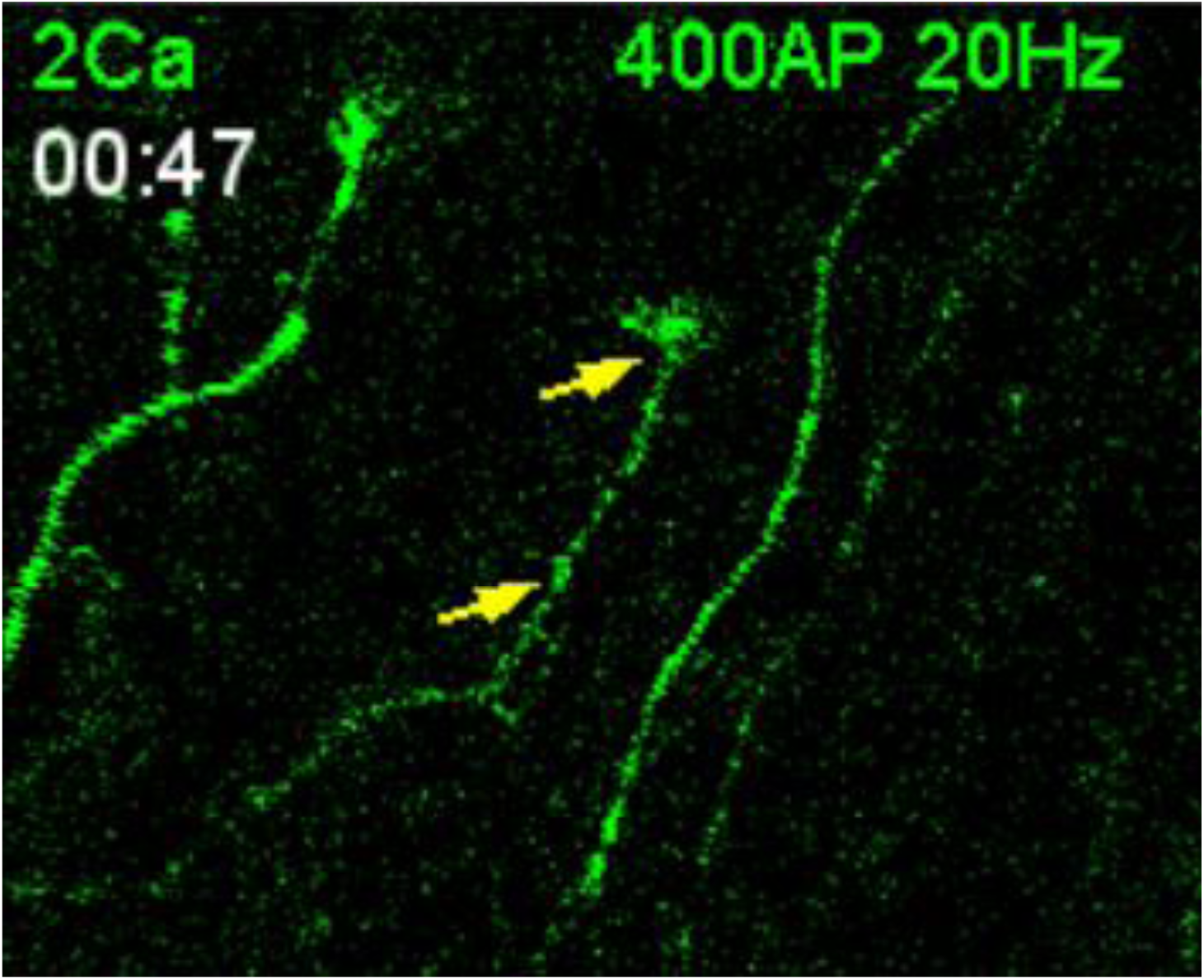
Spy-pHluorin signal from DRG-DH neuron in 2Ca solution.

**Figure 2-video 3.**
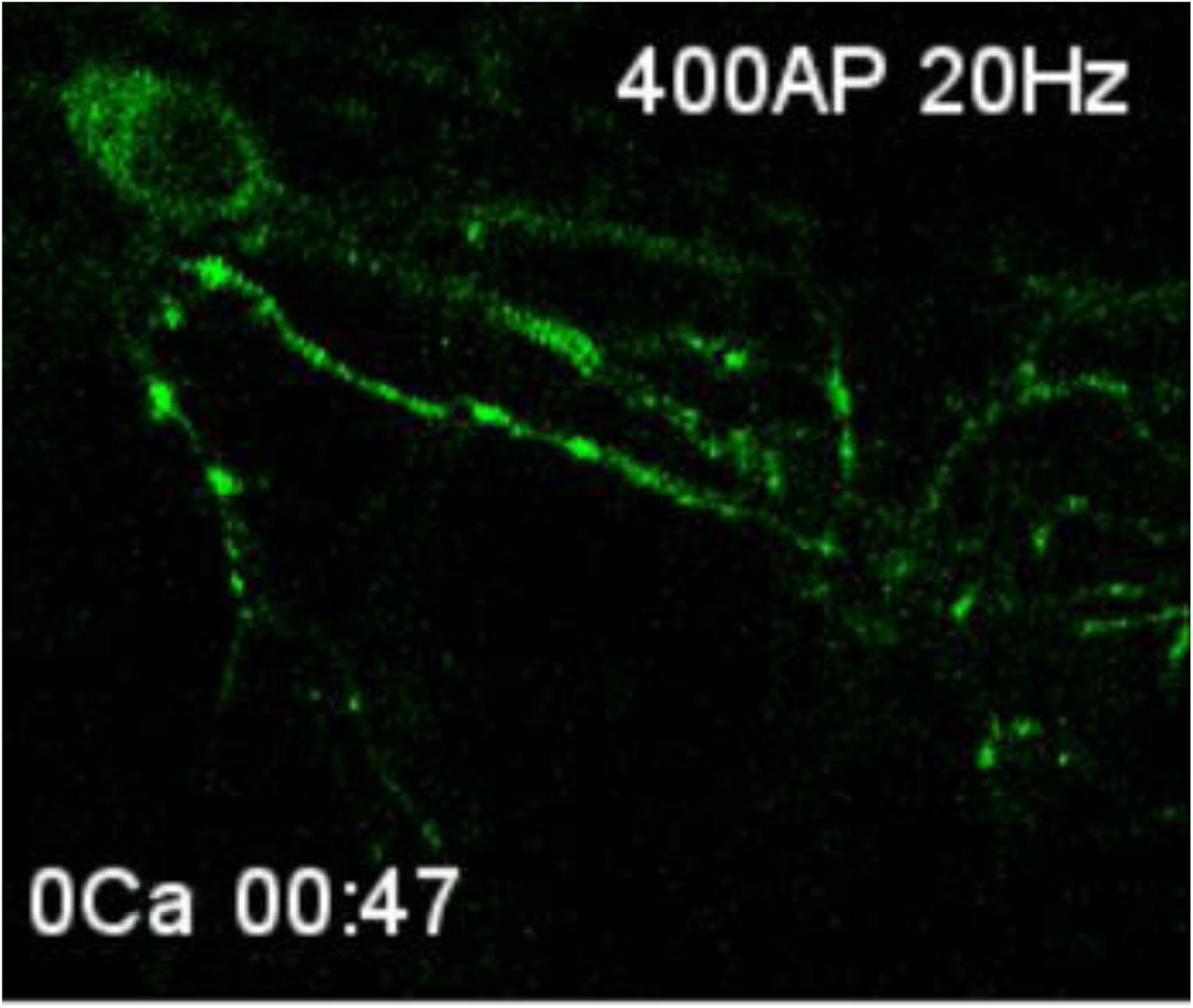
Spy-pHluorin signal from hippocampal neuron in 0Ca solution.

**Figure 2-video 4.**
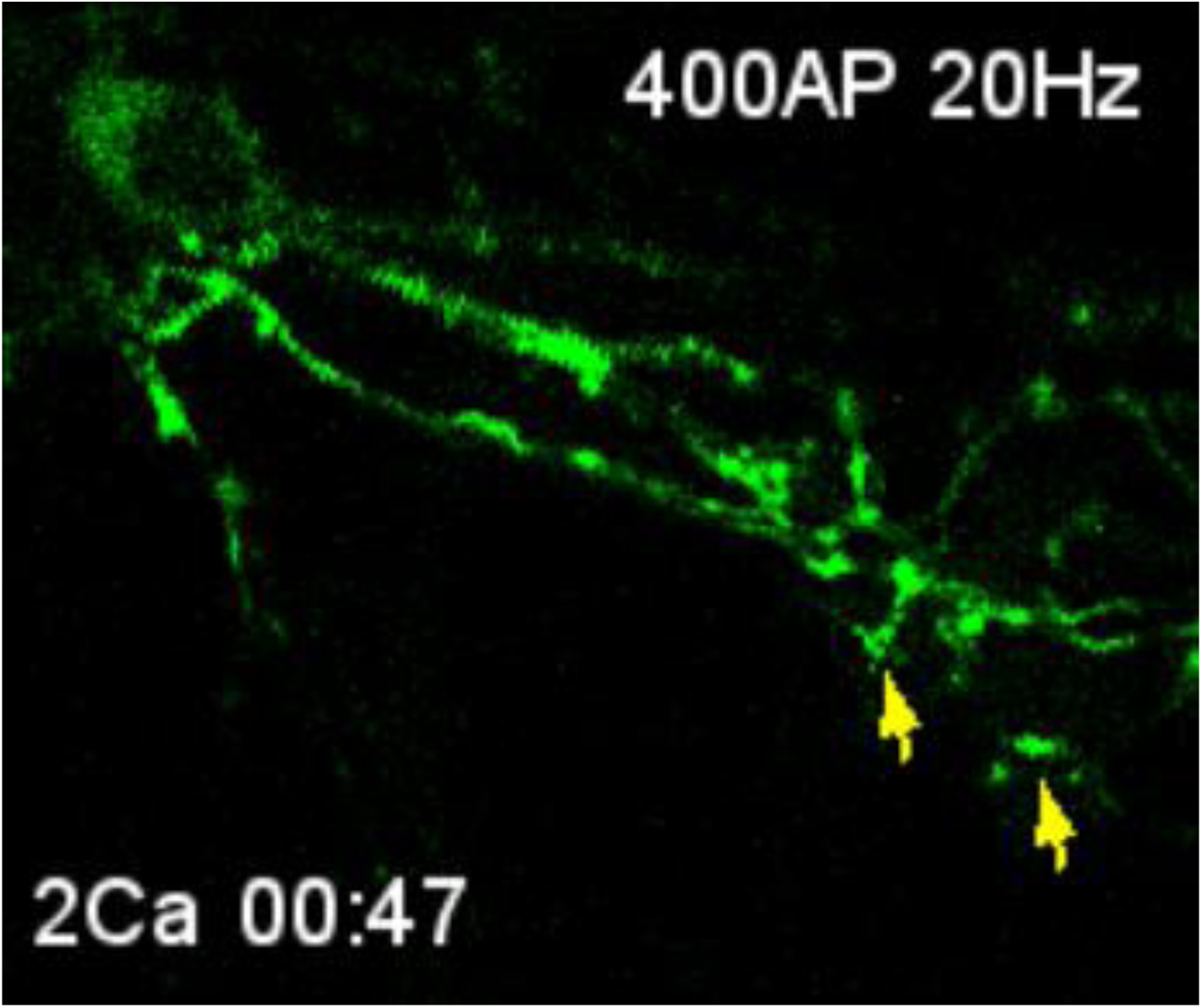
Spy-pHluorin signal from hippocampal neuron in 2Ca solution.

## Notes

### Competing Interest Statement

The authors have declared no competing interest.

